# Cryo-EM structures of the Mycobacterium 50S subunit reveal an intrinsic conformational dynamics of 23S rRNA helices

**DOI:** 10.1101/2022.08.30.505801

**Authors:** Priya Baid, Jayati Sengupta

## Abstract

Pathogenic organisms encounter a broad range of stress conditions within host micro-environment and adopt variety of mechanisms to stall protein translation and protect translational machinery. Structural investigations of the ribosomes isolated from pathogenic and non-pathogenic *Mycobacterium* species have identified several mycobacteria-specific structural features of ribosomal RNA and proteins. Here, we report a growth phase-dependent conformational switch of domain III and IV helices (H54a and H67-H71) of the mycobacterium 23S rRNA. Cryo-electron microscopy (cryo-EM) structures (∼3-4 Å) of the *M. smegmatis* (*Msm*) 50S ribosomal subunit of log-phase manifested that, while H68 possesses the usual stretched conformation in one of the maps, another one exhibits an unprecedented conformation of H68 curling onto a differently oriented H69, indicating an intrinsic dynamic nature of H68. Remarkably, a 2.8Å cryo-EM map of the *Msm* stationary-state 50S subunit unveiled that H68 preferably acquires folded conformation in this state (closely mimicking dormant state). Formation of a bulge-out structure by H68 at the inter-subunit surface of the stationary-state 50S subunit due to the rRNA conformational changes prevents association with 30S subunit and keeps an inactive pool of the 50S subunit representing a ribosome-protection mechanism during dormancy. Evidently, this dynamic nature of H68 is an integral part of the cellular functions of mycobacterium ribosome, and irreversibly arresting H68 flexible motion would stall ribosome function. Thus, this conformational change may be exploited to develop anti-mycobacterium drug molecules.

**Significant statement:** Bacteria utilize several mechanisms to reprogram the protein synthesis machinery so that their metabolism is reduced in the dormant state. Mycobacteria are capable of hiding themselves in a dormant state during physiological stresses. Our study identified a hitherto-unknown folded conformation of the helix 68 (H68) of domain IV of mycobacterial 23S rRNA, which is predominantly present in the stationary state (closely mimicking latency). Our results suggest that this conformational transition is instrumental in keeping an inactive pool of the 50S subunit in the stationary state. Irreversibly arresting such conformational dynamics would lead to protein synthesis shutdown in mycobacteria during dormancy. Thus, this folded conformation of H68 offers an excellent therapeutic intervention site to treat mycobacterial latent infection.

**Highlights:**

- Identification of a hitherto-unknown folded conformation of the helix 68 of mycobacterial 23S rRNA
- H68 conformation transition represents a new ribosome protection mechanism in dormant mycobacteria
- The conformational switch of mycobacterial H68 offers an excellent therapeutic intervention site

## Introduction

The ribosome, one of the most intriguing biomolecular machines (Spirin, 2002), is a ribonucleoprotein complex having an intricate molecular architecture (Schmeing and Ramakrishnan, 2009). Several actinobacterium-specific unique structural features have been identified in mycobacterium 70S ribosome (Kushwaha and Bhushan, 2020). The most striking one is a long (more than 100 nucleotides) helical expansion segment of 23S rRNA, H54a (referred to as ‘Steeple’ (Shasmal and Sengupta, 2012)), that has been identified emerging from the bottom of the L1 stalk side of the 50S (Fig. S1A, B). The tip of H54a reaches at the exit site of mRNA channel (Hentschel et al., 2017; Li et al., 2018b; Shasmal et al., 2016). *Mycobacterium smegmatis* (Msm) 70S ribosome showed that H54a establishes contacts with the small ribosomal subunit protein bS6 to form an intersubunit bridge (Hentschel et al., 2017).

Conformational variability of the H54a has been identified in *Mycobacterium tuberculosis* (*Mtb*) 70S ribosome (Yang et al., 2017). Of note, the ribosome structures of *Mtb* (Cui et al., 2022; Li et al., 2015; Yang et al., 2017) and *Msm* (Hentschel et al., 2017; Li et al., 2018a; Li et al., 2018b; Mishra et al., 2018) are virtually identical (Kushwaha and Bhushan, 2020), suggesting that non-pathogenic *Msm* stands as a good representative of pathogenic *Mtb*, as far as the ribosome structure is concerned. Interestingly, a large conformational change of H54a, as compared to its position in the 70S ribosome, has been identified in 50S ribosomal subunit, where the helix tilts towards the central protuberance (CP) and interacts with the 50S ribosomal protein uL2 and 23S rRNA helix H68 (Li et al., 2015). It is conceivable that species-specific dynamics of the additional ribosomal components play a crucial role in cellular functions of mycobacterial ribosome.

Published structures (Ban et al., 2000; Korostelev et al., 2006; Liu et al., 2017; Watson et al., 2020) of the 70S ribosomes and 50S subunits of different species including mycobacteria, show that H68, part of the domain IV of 23S rRNA, is a long helix (Fig. S1B) occupying the groove behind the L1 protein stalk facing opposite direction from H69 (Ali et al., 2006). Although domain IV of 23S rRNA accounts for a significant portion of the flat intersubunit surface of the large subunit, domain IV helices are interwoven with helices of other domains (Ban et al., 2000) and the helices H68, H69 and H71 are involved in the formation of important intersubunit bridges B2b, B2a and B7a, respectively (Liu and Fredrick, 2016). H67-H71 of domain IV makes a ridge in front of the groove comprising the conserved peptidyl transferase centre (PTC) of domain V (Ban et al., 2000) (Fig. S1B).

In this study, we set out to search for growth phases-specific structural alteration and conformational dynamics of the large ribosomal subunit of *Mycobacterium smegmatis* (*Msm*) using cryo-electron microscopy (cryo-EM). Analysis of high-resolution cryo-EM maps of the *Msm* log phase 50S subunit has led us to identify an altered conformation of H68 in one of the structures. Remarkably, this alternative folded conformation is more prominently present in the large ribosomal subunit of stationary state (closely mimicking latent state) suggesting that the equilibrium of H68 conformational dynamics shifts towards the folded conformation. The molecular model built on a high-resolution cryo-EM map (∼2.8Å) of the *Msm* stationary state 50S subunit revealed that, along with the folding of H68, significant conformational switching occurs in the part of 23S rRNA domain IV comprising H67-H71. Such radical conformational changes in domain IV helices result in formation of a mushroom-head like protruded structure at the intersubunit surface of the 50S subunit and hence aid to prevent 30S subunit association. As a result, active ribosome population for translation is reduced and thereby protein synthesis is down-regulated. Interestingly, density corresponding to the steeple (H54a) is partly visible in this map, away from the fused density of H68-H69, indicating anti-correlated dynamics of H68 and H54a.

Evidently, this dynamic equilibrium of H68 conformations is an integral part of the cellular functions of mycobacterium ribosome, and irreversibly arresting flexible motion of H68 would stall ribosome function suggesting that this conformational change may be exploited to develop anti-mycobacterium drug molecules. Thus, our study not only provides understanding of one of the fundamental mechanisms of mycobacterial ribosome protection and translational down-regulation during stationary/latent state (where the organism exploits intrinsic conformational dynamics of the rRNA), but it also offers a promising drug target for mycobacteria hiding in dormant state.

## Experimental procedures

### Isolation and purification of 70S, 50S and 30S ribosome from *Mycobacterium smegmatis* under log phase and stationary phase of growth

*Mycobacterium smegmatis* (strain MC^2^ 155) cells were grown in MB7H9 medium till log phase (for ∼28 hours) and stationary phase (for ∼ 48 hours) were attained. Cells were harvested by centrifugation (6,000 rpm, 4°C, 10 min.) and washed with Buffer ’A’ (20mM Tris-HCl pH 7.6, 10mM Mg-acetate, 100mM NH_4_Cl, 5mM β-ME). Cells were lysed in Buffer ’A’ by sonicating 10-12 times (amplitude – 90% for 30s with 45s interval), then 50mg/ml lysozyme was added, and cell suspension was incubated at RT for 30 mins, on ice for 20 mins, followed by French Press at 5psi. Cell debris was removed by centrifugation (13,000 rpm, 4°C, 2 hrs.). Supernatant was subjected to ultracentrifugation (28,000 rpm, 4°C, 2 hr. 15mins) in Sorvall AH-629 rotor. Pellet was dissolved in Buffer ’B’ (20mM Tris- HCl pH – 7.6, 10mM Mg-acetate, 30mM NH_4_Cl, 5mM β-ME), salt-washed and homogenized in Buffer ’C’ (20mM Tris- HCl pH – 7.6, 10mMMg-acetate, 1M NH_4_Cl, 5mM β-ME) for 1 hr. Extra protein debris was removed by centrifugation (13,000 rpm, 4°C, 2 hrs.). Supernatant was subjected to ultracentrifugation (28,000 rpm, 4°C, 2 hr. 15min.) in Sorvall AH-629 rotor. Crude ribosome pellet was dissolved in Buffer ’B’ for 70S ribosome purification and in Buffer ’D’ (20mM Tris- HCl pH – 7.6, 0.5mM Mg-acetate, 30mM NH_4_Cl, 5mMβ-ME) for 50S and 30S ribosomal subunits purification. Crude suspension of 70S ribosome was layered on top of sucrose gradient (10%-40% w/v sucrose in Buffer ’B’) to resolve ribosome molecule with ultracentrifugation (28,000 rpm, 4°C, 4 hr. 30 mins), whereas crude ribosome suspension dissolved in Buffer ’D’ was dialyzed against Buffer ’D’ for 6 hr. 30 mins, to allow 70S ribosome to disassociate into 50S and 30S ribosomal subunits. Ribosomal subunits were separated by sucrose density (10%-40% w/v sucrose in Buffer ’D’) gradient ultracentrifugation (28,000 rpm, 4°C, 9hr. 30 mins). Fractions from sucrose gradient were collected using peristaltic pump from bottom to top and checked for the presence of ribosome at A_260_. The fractions corresponding to ribosome peaks (log phase ribosome: L; and stationary phase ribosome: S) were pooled, concentrated and buffer exchanged in a spin concentrator (100,000 MWCO (molecular weight cutoff), Vivaspin) with Buffer ’B’ (High Mg^2+^) for 70S, 50S and 30S. The homogeneity of ribosome was checked on SDS- PAGE and aliquots were stored in −80°C.

### In-Vitro ribosomal subunit re-association Assays

#### (A) Rayligh light scattering studies

Re-association of ribosomal subunits from log (L-50S, L-30S) and stationary phase (S-50S, S-30S) was measured by Rayleigh light scattering in Hitachi F-7000 fluorescence spectrophotometer (with excitation and emission: 2.5 mm slit, wavelength at 350 nm at 90° angle) at 20°C. 50nM of purified 50S from both phases were placed in the cuvette to get the scattering of free 50S subunit. For the re-association reaction, 50nM of purified 50S and 30S subunits from both phases were mixed in ribosome re-association buffer (20 mM Tris-HCl pH 7.6, 10 mM NH_4_Cl, 50 mM KCl, 10 mM Mg-acetate, 5 mM β-ME), and quickly added to the instrument cuvette to get the scattering pattern. The intensity of scattered light was recorded for 5 minutes and 10 sec. The data was plotted in the Origin software and values of starting 10 seconds were omitted while plotting.

#### (B) Sucrose Gradient Density Centrifugation

Two sets of re-association reactions were prepared, one for log phase (L) and another for stationary phase (S). For both sets of reactions, 0.5 μM of 50S was mixed with 0.5 μM of 30S in the ribosome re-association buffer to a final reaction volume of 300 μl and incubated at 37°C for 25 mins, then at RT for 10 mins and finally on ice for 10 mins. After incubation, the reaction mixture was layered on top of 33 ml of sucrose gradient (10%-40% w/v sucrose in ribosome re-association buffer). Ribosomal subunits were separated by sucrose density gradient ultracentrifugation (28,000 rpm, 4°C, 5 hr. 25 mins) in Sorvall AH-629 rotor. Fractions of 500 μl were collected from bottom to top using peristaltic pump. Absorbance at 260 was measured for all the fractions to generate the ribosome profile. Two additional sets of re-association assay were prepared in the same way as mentioned above, with minute variation (one set with L-50S and S-30S and another set with S-50S and L-30S). The peaks corresponding to 70S ribosome, 50S and 30S subunits were pooled separately and concentrated in a spin concentrator (100,000 MWCO (molecular weight cutoff), Vivaspin) and buffer exchanged with Buffer ’B’. The formation of 70S ribosome by in-vitro re-association was checked on SDS- PAGE.

Notably, both of in-vitro ribosomal subunit re-association assay were performed in set of 3 replicates.

### Cryo EM Grid Preparation and data Collection

Cryo-EM grids were prepared with the Vitrobot (FEI) equilibrated at 4°C and 100% relative humidity. *Msm* L-70S ribosome and *Msm* S-70S ribosome - EF-G II complexes were prepared to a final concentration of 150 nM & 200 nM, respectively. 3 μl of samples were loaded on Quantifoil R2/R carbon coated holey grids, which were glow-discharged, and after waiting 25 secs, grids were blotted for 3.5 secs and plunge frozen in liquid ethane.

Preliminary image collection for screening was done using in-house cryo-electron microscope Tecnai G2 Polara equipped with Eagle CCD (data not presented in the manuscript). High-resolution cryo-EM datasets were acquired on a 300-kV FEI Titan Krios electron microscope (at National Center for Biological Science, Bangalore) equipped with CMOS direct detector Falcon III at a calibrated magnification of 101449, yielding a pixel size of 1.38 Å. Defocus values ranged from −1.8 μm to −3.3 μm. Images were acquired in a movie mode using 30 frames/movie (combined dose of 54 electrons/Å^2^).

### Single Particle data processing and generation of 3D structures

Of note, cryo-EM structural studies of 50S subunits were done using ribosome datasets of 70S ribosome containing 50S and 30S subunits as well (in a significant number) to extract 50S particles (both from log and stationary phase, Fig. S2).

#### (A) *Msm* Log-phase ribosome dataset

A total of 1608 movies were acquired from microscope by automated data acquisition using EPU software (FEI). The initial steps of data processing were done in Relion 3 (Zivanov et al., 2018). Movie frames were aligned and combined by dose induced Motion correction using MotionCorr (Zheng et al., 2017). Contrast Transfer Function (CTF) was calculated of motion-corrected movies using CTFFIND 4.1 (Rohou and Grigorieff, 2015). Power spectra and Thon rings for each micrograph was manually analyzed and micrographs with bad CTF were removed. Approximately, 2000 particles were manually picked and extracted with a box size of 350 pixels. Extracted particles were subjected to reference free 2D classification. Good 2D class averages were selected to be used as a template for auto-picking of particles from the entire dataset. A total of 202808 particles were picked by auto-picking, extracted and subjected to multiple rounds of 2D classification to remove bad particles. 3D Initial models were generated from selected good 2D class averages. Initial model of 50S was used as a reference for the first round of 3D classification. The 3D classification resulted in 4 major classes (two 70S, one 50S and one 30S), bad classes were discarded. The class representing 50S was given for multiple rounds of reference-based 3D classification (shown by flow chart in Fig. S2) and resulted in generation of 3 different classes of 50S subunits. All the three classes were subjected to 3D auto refinement. In Relion 3.1 (Zivanov et al., 2018), 3D refined maps were used for particle polishing in train and polish mode. Polished particles were given for CTF refinement (estimation of beamtilt, anisotropic magnification, per particles defocus value and per particle astigmatism). CTF-refined particles were again subjected to 3D auto refinement and another round of particle polishing. Finally, polished particles were used for last 3D auto refinement. The resulting resolution of L1-50S map and L3-50S map were 4.1 Å and 4.8Å, respectively, without post-processing. (Fig. S3) Post processing (auto B-factor sharpening) in Relion 3.1 resulted in a 3.03 Å and 3.22 Å using 0.143 FSC criteria (Rosenthal and Rubinstein, 2015) (Fig. S4). The local resolution estimation of maps was performed using ResMap. The refined maps were sharpened using model based local sharpening tool in PHENIX (Liebschner et al., 2019) for figure preparation.

#### (B) *Msm* Stationary phase ribosome complex dataset

The stationary state 50S subunits particles were isolated from datasets of stationary Phase *Msm* ribosome in complex with *Msm* Elongation Factor-G2 stalled with co-factors. Cloning, expression and purification of Elongation Factor-G2 (EF-G2) from *Mycobacterium smegmatis* were done following standard protocols with slight modifications. The *Msm* ribosome (isolated from stationary state) in complex with EF-G2 stalled with different co-factors were prepared in binding buffer (50mM Tris-HCl; pH-7.8, 70mM NH_4_Cl, 30mM KCl, 7mM Mg-acetate, 1mM DTT). Protein bound ribosome complexes were used for grid preparation.

Two ribosome datasets were used to isolate the stationary state 50S subunits (S-50S). Movies were collected from microscope by automated data acquisition using EPU software (dataset-1; 1719 movies, dataset-2; 1321 movies). The initial steps of data processing were done in Relion 3. Alignment of movie frames and CTF determination were done as described previously. For dataset-1, a few thousand particles were manually picked and extracted with a box size of 350 pixels. Extracted particles were used for reference free 2D class averaging. Good 2D classes have been selected and used as a template for auto picking of particles from entire dataset. A total of 379564 particles were picked and a lot of bad particles were discarded by doing multiple rounds of 2D classification. Good sets of 2D classes were selected and subjected to reference-based 3D classification which resulted in formation of 3 major classes (one 70S and two 50S). Two 50S classes were combined (157711 particles), which were subjected to a series of 2D averaging and 3D classification to get a major class of 50S, which was further subjected to 3D classification. The classification scheme is shown in Fig. S4. *Msm* S-50S particles were also isolated from another MsmS-ribosome-EF-G2 complex dataset. The second S-50S dataset was processed the same way as described before. The structures of the ligand (EF-G2) unbound S-50S 3D classes showed identical features as seen in the unbound 3D classes of S-50S particles extracted from first *Msm* S-ribosome-EF-G2 dataset. Therefore, particles of unbound S-50S classes of these two datasets were merged. Combined 50S particles were subjected to a series of 2D classification to remove some bad particles. Finally, good 2D class averages were subjected to reference-based 3D classification (shown in Fig. S8). The final refined map consisted of 172451 particles, which was processed further for particles polishing, CTF refinement and 3D auto refinement as described above (Fig. S8). The resolution of final S-50S map was 2.8 Å after post-processing (auto B-factor sharpening) in Relion 3.1 and local resolution estimation was done using ResMap. The Final 3D refined map was sharpened with model based local sharpening tool in PHENIX for figure preparation (Fig. S9).

### Atomic model building, refinement and validation

All Cryo-EM maps were initially analysed in UCSF ChimeraX by thorough segmentations to check the presence of ribosomal RNA and all ribosomal proteins (Figs. S5, S6 and S10). The model coordinates of *Msm* 50S (PDB ID- 5O60) has been used as reference for model building and fitting. Model coordinates were initially aligned and fitted with cryo-EM map using Chimera. Further, manual fitting of steeple (H5Aa), H67-H71 and L2 of L1-50S and S-50S was done in Coot using refine and regularize mode (Fig S5E-G & Fig S10E-G). Notably, backbone trace model fitting was done for H68-69 region in S-50S, as this region was comparative low resolved (Fig. S11). Models were subjected to real-space refinement in Phenix after crude fitting in Coot (Emsley et al., 2010), then iteratively fitted and refined in Coot and Phenix, respectively, until final model had been generated. Validation of model coordinates was done (Table 1) by comprehensive validation in Phenix using MolProbity (Chen et al., 2010). Illustrations were prepared with UCSF Chimera (Pettersen et al., 2004), UCSF ChimeraX (https://www.rbvi.ucsf.edu/chimerax/) and PyMOL (Schrödinger, LLC.).

**Table 1.**
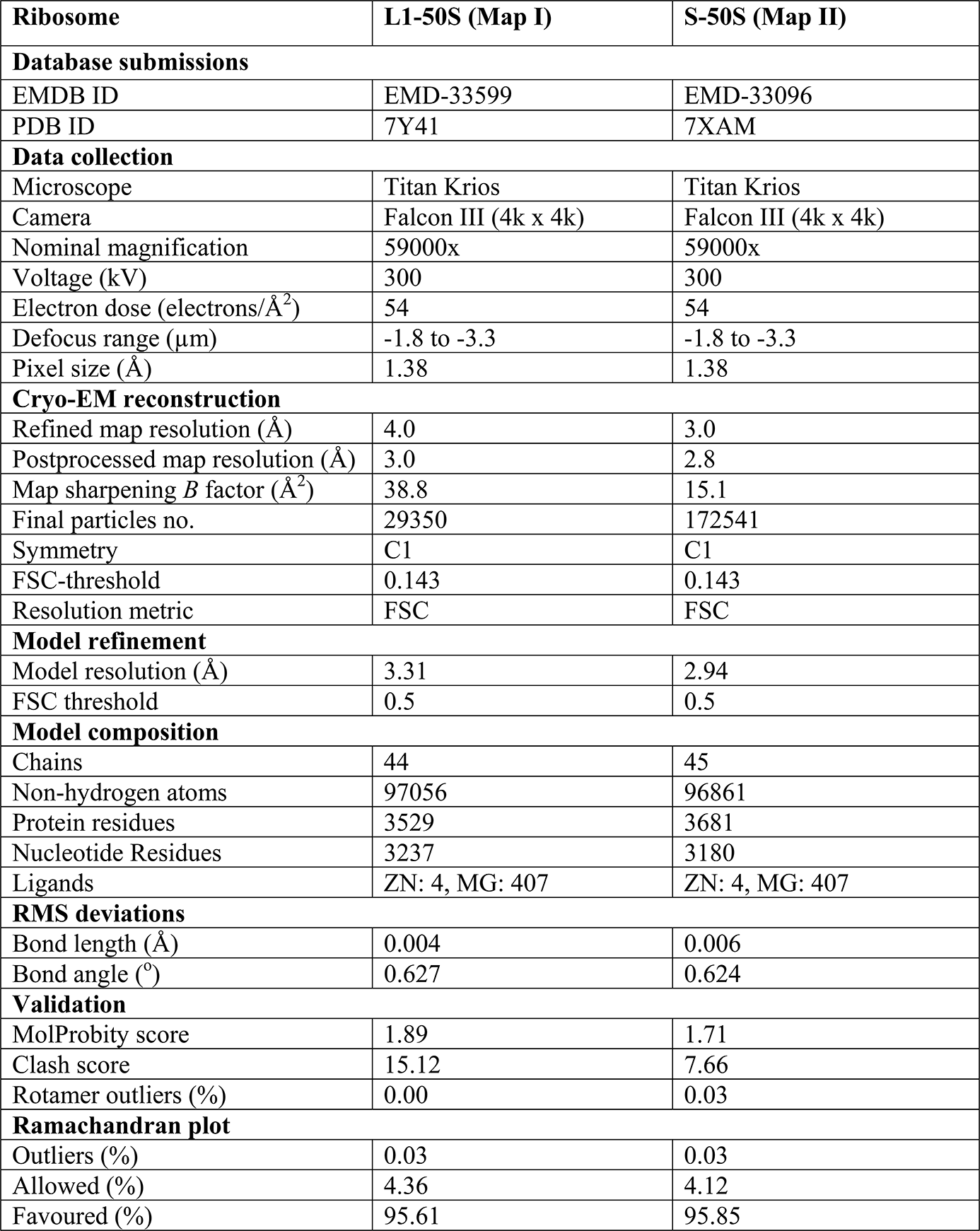
Data collection, refinement and validation statistics.

### 3D structure prediction of H67-H71 of Mycobacterial 23S rRNA

The rRNA sequence of H67-H71 of *M. smegmatis and M. tuberculosis* were obtained from NCBI, USA. The sequence was submitted to RNAComposer (an automated RNA structure 3D modelling server) for 3D prediction, where ContextFold was used as a method of choice for secondary structure prediction.

## Results and discussion

### Intrinsic dynamic nature of helix 68 of the mycobacterium 23S rRNA

*Msm* ribosome sample, purified from exponential phase of growth, was used for cryo-EM structure determination. A pool of 50S particles were extracted from a ribosome image dataset containing 70S ribosome as well as the subunits (Fig. S2) using multiple rounds of 2D classifications, followed by 3D classification (Fig. S3). Further, extensive 3D classification and refinement of extracted 50S particles resulted in two major reconstructions of *Msm* 50S subunit, termed as map L1-50S (Fig. 1A) and L3-50S (Fig. 1B) (res 3.03Å and 3.22Å post b-factor sharpening, respectively (Fig. S4A-B, E-F), along with another minor class having a small population of particles (map L2-50S, ∼18Å resolution; Fig. S3). Although, the core structures of both maps are highly resolved, the resolution dips towards peripheral regions as visualized by local resolution estimation of refined reconstructions (Fig. S4D, H). Segmentation of the density maps showed occupancy of almost all the ribosomal proteins except bL9 and uL10 in L1-50S and L3-50S. Densities corresponding to 5S rRNA and 23S rRNA were also distinctly visible (Fig S5A-D, S6).

**Figure 1:**
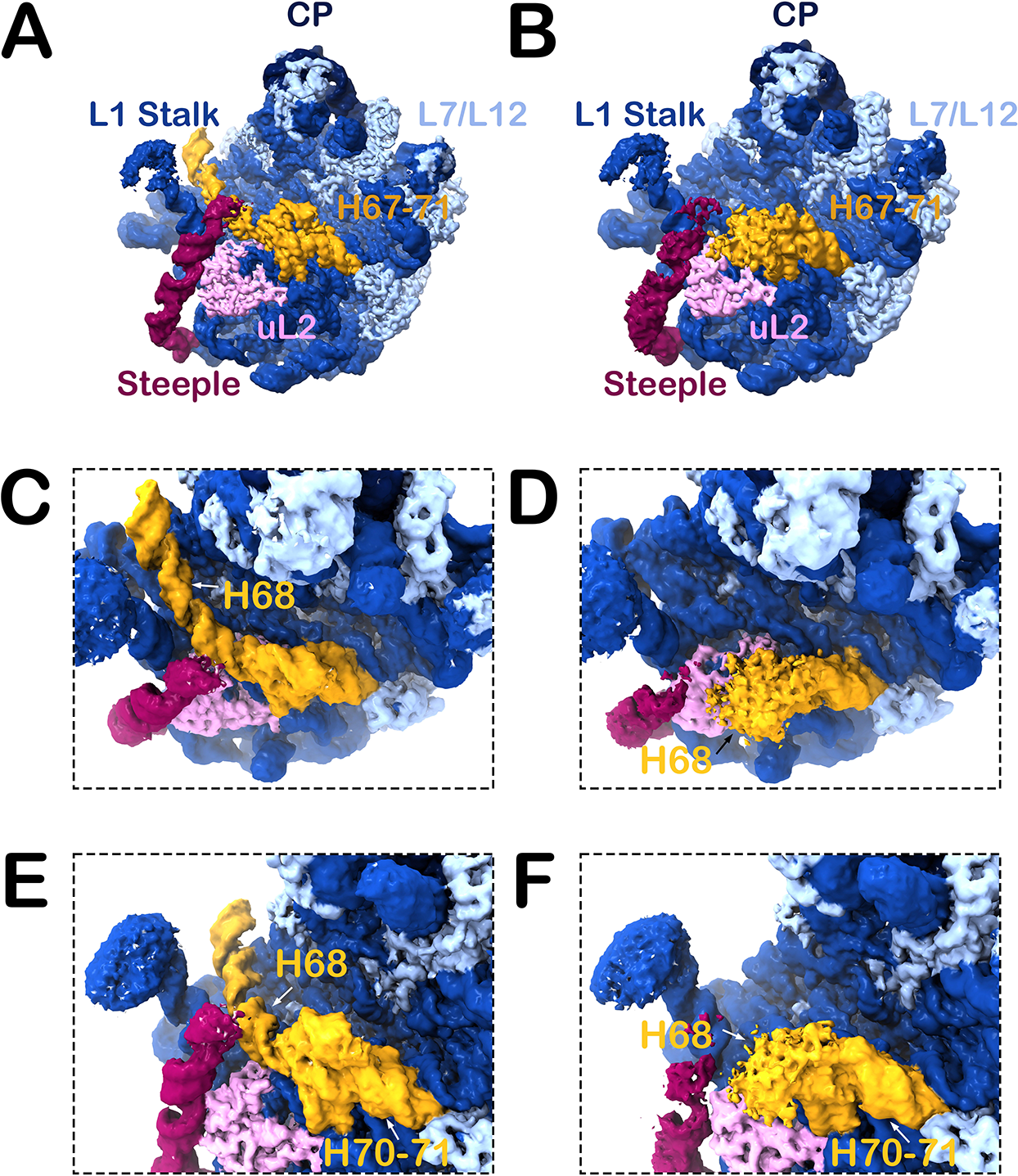
Conformational dynamics of Helix 68 of the log-phase *Msm* 50S subunit. (A & B) Intersubunit view of cryo-EM maps of L1-50S and L3-50S, respectively. Densities of rRNA and ribosomal protein are color-coded (23S rRNA, dark blue; 5S rRNA, blackish blue; r-proteins, light blue; H67-H71, mustard yellow; Steeple, magenta; and uL2, pink). H68 lies in elongated position in L1-50S (A), as observed previously, and attains folded conformation in L3-50S (B). (C-F) Two different close-up views are presented showing H68 of L1-50S in the straight position (C & E), and H68 of L3-50S in an alternative curled conformation (D&F). The steeple (H54a) of the 23S rRNA is seen in an inward tilted position for both the maps. Landmarks for the 50S subunit: CP, central protuberance; L7/L12, stalk made by L7/L12 proteins; L1, L1 protein stalk.

The overall architecture of 23S rRNA in the L1-50S is very similar to the available structures of mycobacterial ribosome (Fig. 1C, E). In contrast, an unusual conformational change in the H67-H71 region was observed in L3-50S (Fig. 1B), where H68 attains a compact mushroom head-like bulge-out feature on the H67 stalk (Fig. 1D, F). The bulge-out structure at the intersubunit surface of L3-50S (Fig. 1D, F) likely prevent its association with the 30S subunit, and thus L3-50S needs a conformational switch to L1-50S, which is compatible to form the active 70S ribosome. Clearly, a dynamic equilibrium exists in H67-H71 conformations of the log phase 50S subunits. Interestingly, dynamic nature of H68 has also been reported in recent study of *S. aureus* 50S subunit, where a ∼70° displacement of H68 from its usual position towards L1 stalk is seen (Cimicata et al., 2022). Notably, 3D structures prediction of mycobacterial H67-H71 by RNAComposer web server, resulted in a curved conformation of H68 of mycobacterium 23S rRNA (Fig. S1D-E), likely indicating an intrinsic tendency of H68 to fold. H54a (Steeple) stays in a tilted position in both the maps, as observed in *Mtb* large subunit structure (Yang et al., 2017), bringing it close to CP (Fig. 1). Notably, map L2-50S shows (Fig. S3) a folded conformation of H68 (similar to map L3-50S) and H54a in an outward position (similar to its position in the previously published *Msm 70S* ribosome structures).

### Growth phase dependent changes in the mycobacterium ribosome population

In order to understand whether any growth phase dependent differences in the association of 50S and 30S subunits to form the 70S ribosome, ribosomal subunits of *M. smegmatis* were purified from exponential and stationary phases of growth using sucrose density gradient separation (see Materials and Methods). Purified 50S and 30S subunits from both the phases were used to perform *in-vitro* re-association assay by sucrose density gradient centrifugation and Rayligh light scattering (excitation and emission at 350 nm) to monitor the formation of 70S ribosome.

Interestingly, analysis of sucrose density gradient ribosome profile of subunit re-association showed significant reduction in 70S ribosome formation by the 30S and 50S subunits purified from stationary phase (S-30S, S50S) as compared to that of the log phase (L-30S, L50S) (Fig. 2A, B). Rayleigh light scattering analysis also showed that stationary-state 30S and 50S subunits do not associate as efficiently as the log-phase 30S and 50S subunits do (Fig. 2C, D). Further, re-association assay by sucrose density gradient ribosome profiling showed that association of S-50S and L-30S gets hindered, while L-50S and S-30S subunit efficiently associates to form 70S ribosome (Fig. 2E, F), suggested that 50S subunits, not the 30S subunits, of the stationary phase are responsible for inefficient re-association.

**Figure 2:**
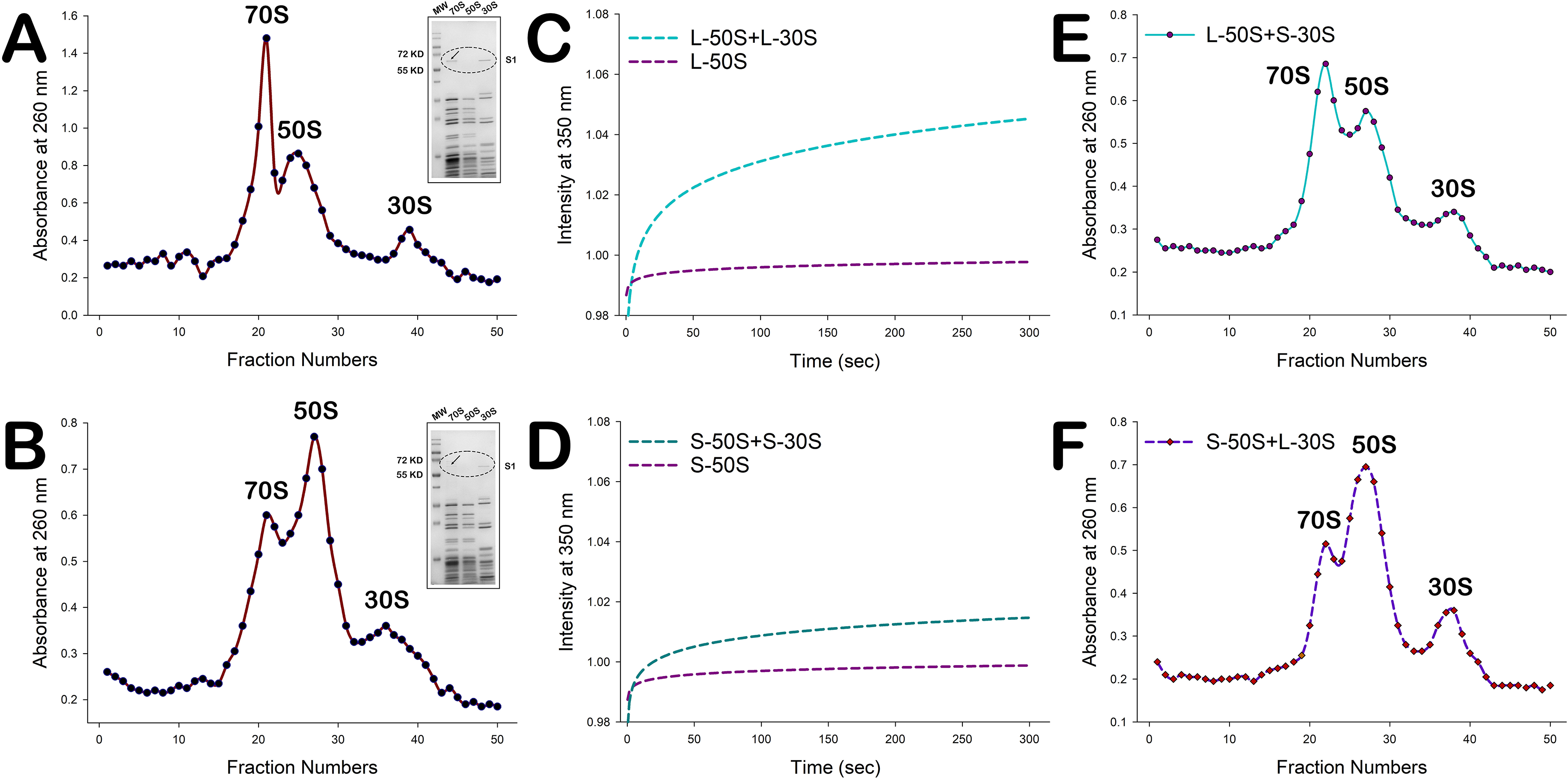
*In-vitro* re-association study of *Msm* ribosomal subunits from log and stationary phases. (A-D) Sucrose density gradient (SDG) profile and light scattering (LS) analysis of ribosomal subunit association from log phase (A & C) and stationary phase (B &D) are shown. SDG profiling and LS analysis shows formation of 70S to a significant level by re-association of subunits purified from log phase, whereas, 70S ribosome formation is substantially reduced in stationary phase subunits. SDS-PAGE analysis of ribosome peaks of SDG profile shown in insets of (A & B). (E) SDG profile of re-association of L-50S and S-30S and (F) SDG profile of re-association of S-50S and L-30S show that 70S formation is comparatively reduced in the presence of S-50S. All the re-association assay were repeated 3 times. One of the representative results is shown here.

A comparative analysis of ribosome profiles of purified 70S ribosome from *Msm* log and stationary phases clearly showed that occurrence of the 50S subunit peak along with 70S ribosome is higher in stationary phase as compared to log phase (Fig. S7Ai, Bi). Moreover, when stored *Msm*70S ribosome purified from log and stationary phases were again loaded onto sucrose density gradient, only a single 70S ribosome peak was observed in log phase ribosome profile. In contrast a clearly detectable peak of the 50S subunit was also observed along with the 70S ribosome peak (Fig. S7Aii, Bii), indicating that the equilibrium between 70S ribosome and its subunits tends to shift towards ribosomal subunits in stationary phase. A plausible explanation for this observation would be that dissociated subunits are unable to efficiently re-associate to form active 70S ribosome due to some structural changes, creating a pool of inactive 50S subunits in the stationary phase. Since alternative conformation of H67-H71 in the 23S rRNA would inhibit 70S ribosome formation, we were curious to know if this conformation of H67-H71 is more prominently present in the stationary phase 50S ribosomal subunit.

### Helix 68 preferably adopts folded conformation in the stationary state Msm50S

In order to gain further structural insights into the domain IV conformation of the stationary state 50S ribosomal subunit (S-50S), we carried out cryo-EM 3D reconstruction of the S-50S following the same workflow as applied for structural analysis of the log phase 50S subunit (Fig. S2). S-50S population was extracted from two datasets (see Materials and methods) of *M. smegmatis* ribosome in complex with elongation factor G2 (EF-G2, an ortholog of EF-G found in mycobacteria), where ribosomes were isolated from the stationary phase of growth. Repeated 2D and 3D classifications of extracted 50S subunit particles from both the datasets were performed and analysis of 3D classes showed that in one dataset, out of 5 major classes, 4 classes of ligand free and one class of EF-G2-bound 50S subunit were generated, while other dataset attained all ligand-free 3D classes (not shown). Remarkably, H68 was found in folded form in the EF-G2-bound class as well as in all the ligand-free classes of the S-50S (Fig. S8), suggesting that the folding of H68 is an intrinsic property of the stationary phase 50S subunit and not induced by EF-G2 binding.

Notably, features of all 3D classes of ligand-free 50S subunits were identical from both the datasets, therefore particles representing these classes were merged, polished and 3D reconstructed to generate a high resolution cryo-EM structure of stationary phase 50S subunit (map S-50S), which reached an average ∼2.8Å resolution after b-factor sharpening (post-processing) (Fig. 3A, Fig. S9 A, D). Local resolution estimation of density map clearly shows that the core region of 50S subunit has been resolved to sub-3Å resolution, while peripheral regions such as elements at intersubunit surface, central protuberance, L1 stalk and L7/L12 are comparatively less resolved (∼4-5.5Å) (Fig. S9C). Full densities attributable to all the ribosomal proteins including bL9, 5S rRNA and 23S rRNA are distinctly visible in segmental exploration of sharpened map (Fig. S10A-D). Closer inspection of S-50S map revealed that H67-H71 of domain IV has undergone more prominent conformational alteration, similar to as observed in L3-50S, implying role of this significant conformational change of H67-H71 in high anti-association property of the S-50S (Fig. 3B). Interestingly, the density corresponding to the Steeple (H54a) is partially visible and has an outward placement (Fig. 3C) as compared to its density in L1-50S and L3-50S structures, where full density corresponding to the Steeple is visible (tilted towards the CP), showing its dynamic nature.

**Figure 3:**
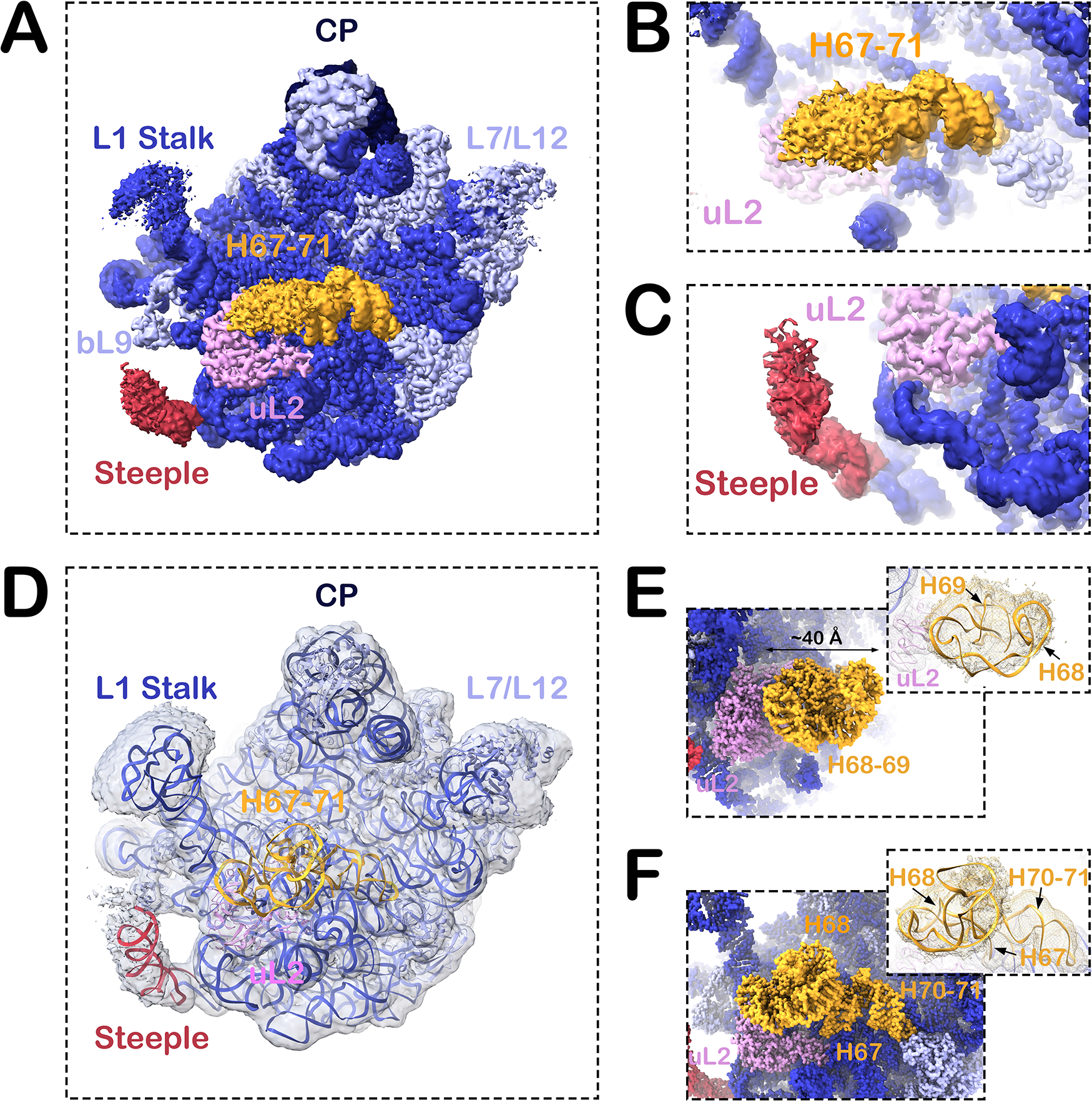
Structural analysis of S*-*50S showing rotated conformation of H68. (A) Intersubunit view of 2.8Å cryo-EM density map of the *Msm* stationary phase 50S subunit. Densities of rRNA and ribosomal protein are color-coded (23S rRNA, dark blue; 5S rRNA, blackish blue; r-proteins, light blue; H67-H71, mustard yellow; Steeple, red; and uL2, pink). (B & C) Close-up views of H67-H71 and the Steeple, showing H68 in rotated conformation (B), and partial density of the Steeple in outward position (C). Weaker density corresponding to H68 suggests dynamic nature of the folded conformation. (D) Fitted molecular coordinates of S-50S in the refined density map (semitransparent blue). (E & F) Close-up views of the molecular models of the rotated H68 are shown in different orientations. Fitting of the molecular coordinates into the density map are shown in respective insets. Landmarks are the same as Figure 1.

Visualization of the H67-H71 region in the refined (prior to b-factor sharpening) and sharpened maps of S-50S (Fig. S11) revealed the emergence of a mushroom head-like bulge out density of H68-H69 in the refined map while moving from high to low contour levels (density appears fragmented in the sharpened map even at lower contour level (Fig. S11)). The densities corresponding to H67 and H70-H71 on the other hand were clearly visible at high contour level even in the sharpened map (Fig. S11). Although it is clear that H68 prefers to adopt rotated conformation in S-50S, comparatively lower resolution of the fused density of H68-H69 indicates conformational flexibility of this region. The refined map allowed us to construct backbone trace model of H68-H69 which could be satisfactorily accommodated into the bulge-out density (Fig 3D).

A close inspection of the fitted molecular model of H67-H71 region revealed that H68 folds itself in twirling conformation over H67 and, along with ∼90° rotated H69, acquires a mushroom head-like configuration (>40 Å bulging out) (Fig. 3E). H70-H71 in an altered conformation acts as an additional support to this bulge out structure (Fig. 3F).

Taken together, the above observation suggested that the conformational switch of H67-H71 region would hinder formation of important intersubunit bridges between 50S and 30S subunits (Fig. S1B) and thus association of ribosomal subunits would be prevented in the stationary state (closely mimicking latent state).

### Molecular mechanisms underlying the conformational transition of H67-H71

The cryo-EM structure of *Msm* L1-50S in compliance with previously reported structure of *Mtb* 50S subunit shows that elongated structure of H68 and partly unstructured H70-71 of domain IV establishes some inter and intra-domain molecular interactions with different rRNA helices (Fig. 1A). Closer analysis of fitted model coordinates of *Msm* L1-50S reveal details of all such stabilizing interactions, as visualized H68 creates stable molecular interaction with H22-H88 of domain I-V, H54a (steeple) of domain III, H66 of domain IV and with H75 of domain V (Fig. 4A-D), further electrostatic interactions are also observed between some basic amino acids (H245 & R256) of bL2 with RNA backbone of H68 (Fig. 4E-F). Evidently, A2162 of H70 makes a base stacking interaction with A2003 of H65 of domain IV (Fig. 4G), while U2168 and U2179 interacts with H92 of domain V (A-loop) (Fig. 4H), further U2163 and A2190 of unstructured part of H70 build connection with H93 of domain V (Fig. 4I-J, Fig. S12A-C). Remarkably, all these inter-molecular interactions of H68 and H70-71 are disrupted due to the drastic conformational transition of H67-71 observed in L3-50S and S-50S.

**Figure 4:**
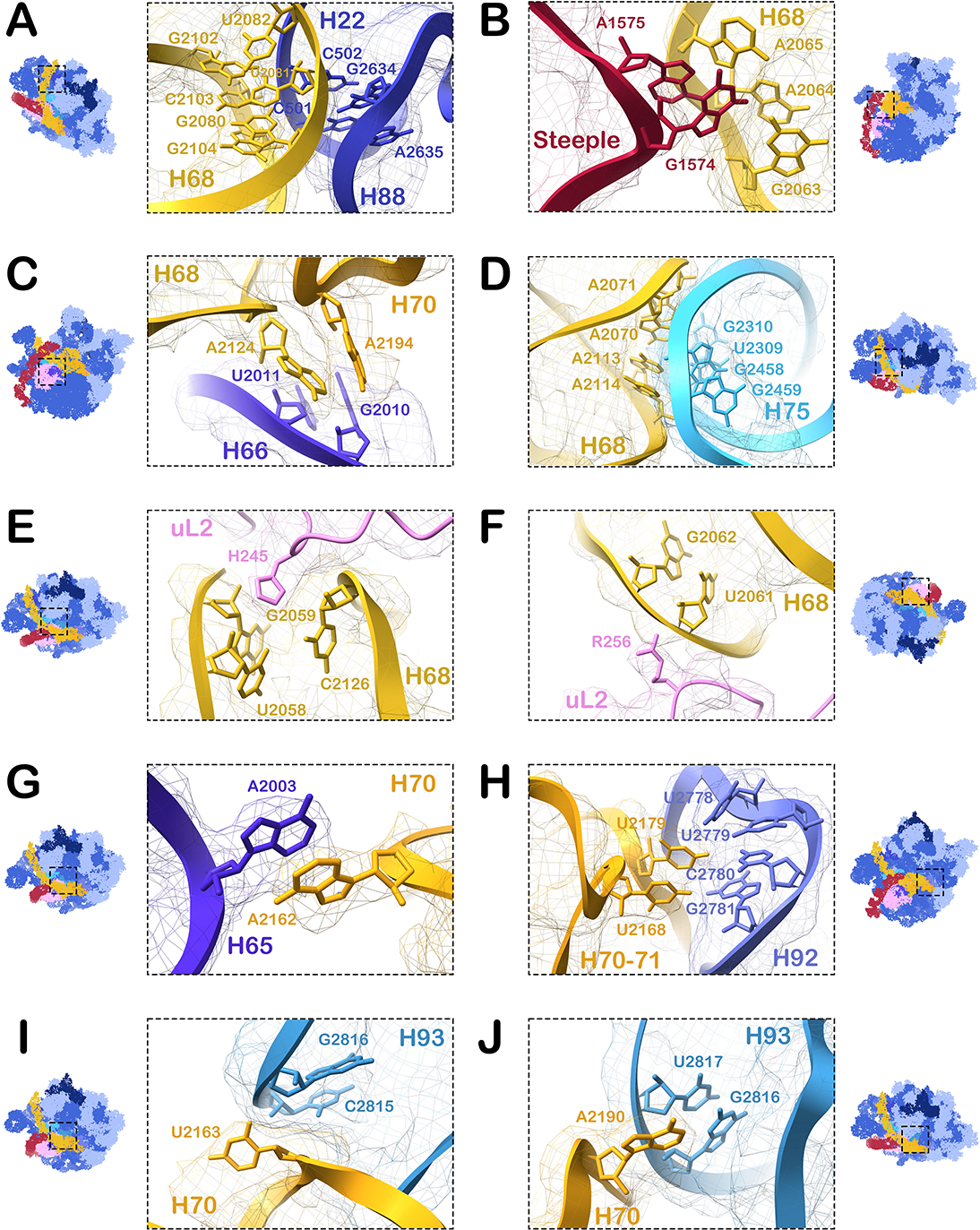
Interactions of H68 and H70-H71 with the ribosomal components in L1-50S. (A-F) Elongated conformation of H68 in L1-50S interacts with H22 and H88 of domain I and V, respectively (A), H54a (steeple) of domain III (B), H66 of domain IV (C), H75 of domain V (D), and with the ribosomal protein uL2 (E & F). (G-J) Partly unstructured H70-H71 of L1-50S interacts with H65 of domain IV (G), H92 of domain V (H), and H93 of domain V (I & J). All of these interactions are disrupted upon conformational switching of H67-H71 in stationary phase 50S subunit.

Comparative analysis of the H67-H71 of domain IV in molecular models of L1-50S and S-50S unveils the movement of H68 from extended to folded conformation (Fig. 5A, Movie S1) and associated conformational alteration of H67, H69 and H71 (Fig. 5B-D). H67 slightly tilts (∼5-6 Å) towards intersubunit surface in S-50S as compared to its position in L1-50S (Fig. 5B, Movie S2). Furthermore, coordinating with the movement of H67, H70-71 undergoes intra-helical molecular rearrangement to attain more structured helical configuration (Fig. 5C, Movie S3) following detachment from H-93 (Fig. S12D-F) in S-50S. Interestingly, H69 also moves laterally to attain a nearly 90°-flipped position in S-50S (Fig. 5D, Movie S4). Similar conformational repositioning of H69 has been reported in pre-60S subunit structure of a yeast ribosome (Wu et al., 2016). Altogether, these tiny changes in the conformations of H67, H69 and H71 bring out a gross conformational change in H68 as bacteria shifts from log to stationary phase.

**Figure 5:**
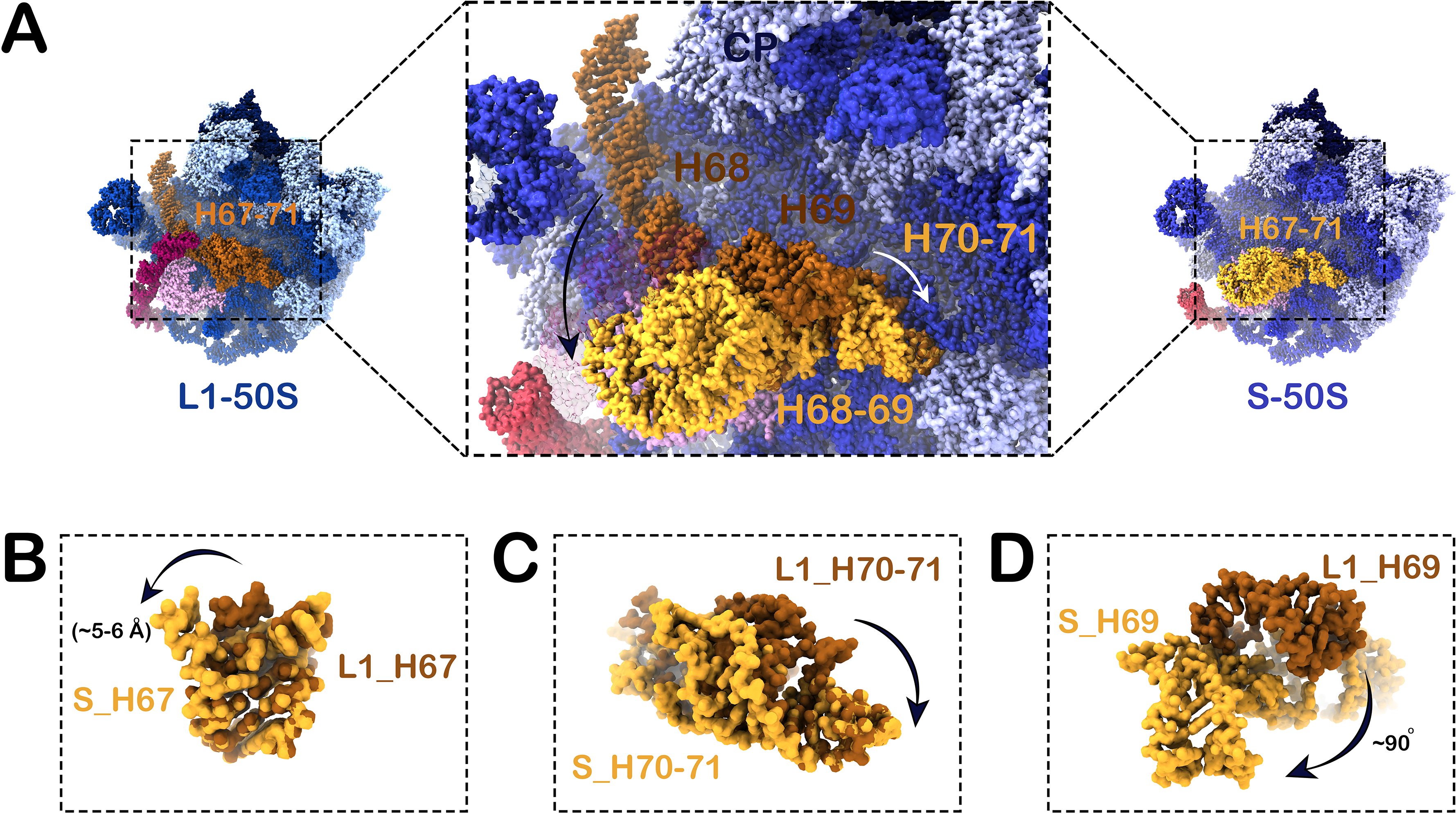
Structural differences of the H67-H71 in L1-50S and S-50S. (A) Close-up view (middle) of the conformational transition of H68 from elongated to folded structures is presented (super-imposed molecular models of H67-H71) from L1-50S and S-50S (thumbnails are shown in two sides). (B-D) Comparisons of H67, H70-71, and H69 from L1-50S and S-50S are shown. As visualised, H67 protrudes ∼5-6 Å towards intersubunit space in S-50S (B), H70-71 becomes more structured after intra-helical molecular rearrangement in S-50S (C), and H69 flips ∼90° from its position in L1-50S to S-50S (D).

### Role of ribosomal protein uL2 in regulating H67-H71 dynamics

Analysis of L1-50S and S-50S structures showed that uL2 is the only ribosomal protein to be present in the close vicinity of H67-H71 of 23SrRNA domain IV (Fig. 1A, Fig. 3A). As mentioned above, uL2 creates electrostatic interactions with H68 in L1-50S (Fig. 4E-F) and further examination of uL2 in L1-50S showed interaction of amino acid residues G238-T240 to minor groove of H93 (nucleotide residues 2814-2815) (Fig. 6A), while S241 of uL2 forms connection with U2195 of H70 (Fig. 6B). In compliance with previously reported study on *Mtb* 50S subunit, we have also observed interactions between three loops of solvent exposed side of uL2 (amino acid residue D169, R134 and N122) with minor groove of H54a (nucleotide residue C1562, C1561 and U1612) in L1-50S (Fig. 6C-E).

**Figure 6:**
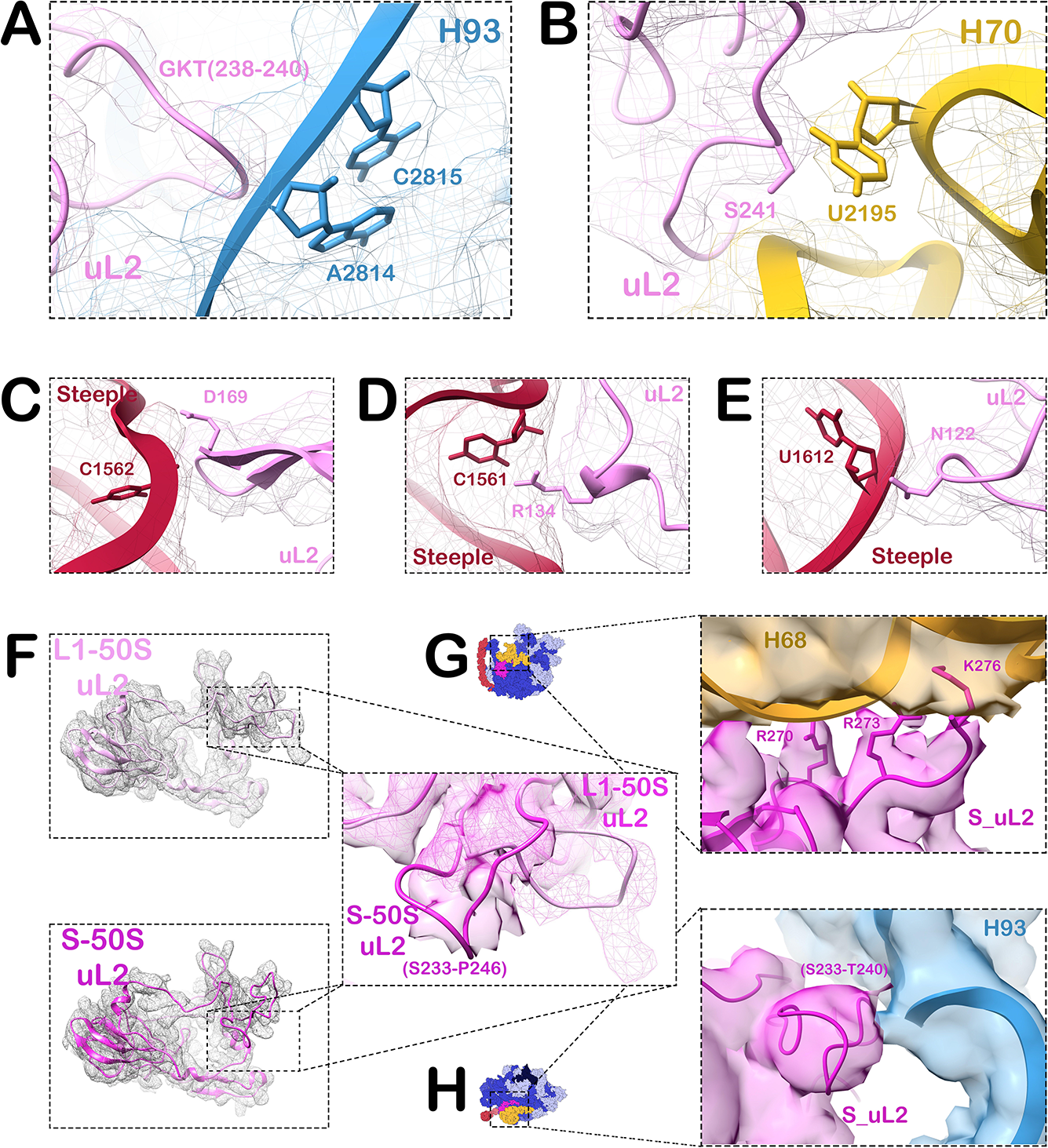
Growth phase dependent conformational switch of ribosomal protein uL2. (A-E) Molecular interactions made by ribosomal protein uL2 with rRNA helices as seen in L1-50S. The protein uL2 interacts with H93 of domain V (A), H70 of domain IV (B) and the Steeple (H54a) of domain III (C-E) in L1-50S. (F) Conformational rearrangement in an unstructured loop (S233-P246) of uL2 from log to stationary phase is seen in close-up view of super-imposed isolated density of uL2 from L1-50S and S-50S. (G & H) Close-up views of interactions of uL2 with H93 and H68 in S-50S shows that rearranged loop of uL2 (S233-T240) interacts with H93 (G) and positively charged amino acid at C-terminal end of uL2 makes connection with folded conformation of H68 (H), suggesting structural changes in uL2 stabilizes folded conformation of H68 in S-50S.

Furthermore, to decipher growth phase-specific structural changes in uL2 that accompanies with the conformational transition of H67-H71, we have inspected segmented density along with the fitted atomic model of uL2 from L1-50S and S-50S. An unstructured loop (made up of amino acid residues S233-P246) towards C-terminal end of protein uL2 moves laterally to gain a tiny bulging shape in S-50S as observed in superimposed models of L1-50S and S-50S (Fig. 6F). Remarkably, above mentioned connections of uL2 with H68 and H70 of domain IV gets disrupted, while interaction between uL2 (amino acid residues S233-T240) and H93 of domain V gets rearranged during repositioning of unstructured loop from L1-50S to S-50S (Fig. 6G). Further, positively charged amino acid residues (R270, R273 and K276) at C-terminal end of uL2, establish interaction with rotated configuration of H68 (Fig. 6H). Evidently, in coordination with conformational transition of H67-H71, structural rearrangements in uL2 during stationary phase support the folded conformation of H68 in S-50S.

### Anti-correlated, coordinated movement of H68 and the Steeple

Previously, we have predicted a dynamic nature of the Steeple based on its free-standing structure in *Msm*70S ribosome (Shasmal and Sengupta, 2012). Reported structure of *Mtb* 50S subunit showed a tilted position of H54a (Steeple) towards central protuberance (CP) of the 50S subunit (Li et al., 2015). Interestingly, identical conformation of H54a was observed in *Msm* L1-50S, where H54a bends towards CP and H68 remains in elongated conformation (Fig. 1A). Inspection of the density maps of *Msm* 50S subunit from different growth phases revealed coordinated positional alterations of the Steeple (H54a) with the conformational switch of H68 (Fig. 1A-B, Fig 3A).

Notably, interactions with the elongated structure of H68 and uL2 stabilize H54a in a tilted position in L1-50S (Fig. 4B, Fig. 6C-E). Although H54a stays in tilted position in L3-50S, weaker density corresponding to H54a in this map suggests its conformational variability. Upon H68 conformational transition from elongated to rotated structure in L3-50S, the H68-H54a interaction gets disrupted (Fig.1B) which apparently destabilizes H54a. Furthermore in S-50S, rotated conformation of H68, along with the conformational rearrangement of uL2, results in an outward motion of the Steeple (Fig. 3A, C). Evidently, to stabilize the tilted position, H54a needs to anchor onto the elongated H68 in the 50S subunit. Comparison of molecular model of L1-50S, L3-50S and S-50S indicated that L3-50S likely represents an intermediate conformation of L1-50S and S-50S (Fig. S13). It may be note here, the density map of the minor class generated for the log phase 50S subunit (L2-50S; Fig. S3) resembles the structural features of S-50S.

The above observations, in compliance with our structural analysis of different conformations of the 50S subunit (L1-50S, L3-50S and S-50S), has led us to propose that coordinated movements of H54a and H68 (the outward-inward motion of H54a and the curling motion of H68) are anti-correlated. In other words, conformational transition of H68 from elongated to folded form is coupled with the outward movement of tilted H54a (Movie S5).

Taken together, we suggests that dynamic equilibrium exists among different conformations (L1-, L2- and L3-50S) of the *Msm* 50S subunit during log phase and the equilibrium shifts to S-50S conformation as bacteria enters stationary phase of growth, implying that the anti-association nature of the S-50S conformation inhibits binding with *Msm* 30S subunit and is utilized in translational regulation in order to accomplish restricted protein synthesis in stationary phase.

### Conclusion

*Mycobacterium tuberculosis* (*Mtb*), an opportunistic microorganism, is known to hide itself in a dormant state under unfavorable physiological conditions. Bacteria reprogram the protein synthesis machinery to abate their metabolism in the latent state. Several energy conservation mechanisms in bacteria are known where different non-canonical factors are involved in the translation regulation through formation of inactive 70S ribosome, or 100S ribosome dimers (Starosta et al., 2014). Interestingly, 100S formation is not a very common phenomenon in mycobacterium (Kushwaha and Bhushan, 2020).

Our structural study of *M. smegmatis* large ribosomal subunit revealed a growth phase dependent hitherto unknown intrinsic conformational dynamics of H67-H71 of domain IV of mycobacterial 23S rRNA and this radical conformational transition is likely instrumental in keeping an inactive pool of the 50S subunit in the stationary state. The unusual conformational alteration of H67-H71 region, prominently present in the stationary state, creates a mushroom head-like protrusion at the intersubunit surface of the 50S subunit and obstructs formation of several important intersubunit bridges B2a, B2b and B7a between ribosomal subunits and thereby prevents active ribosome formation in the stationary state (represents a state closely mimicking dormancy) (Fig. 7). Thus, our study has unveiled a novel mechanism of ribosome protection during dormancy in *Mycobacterium*.

**Figure 7:**
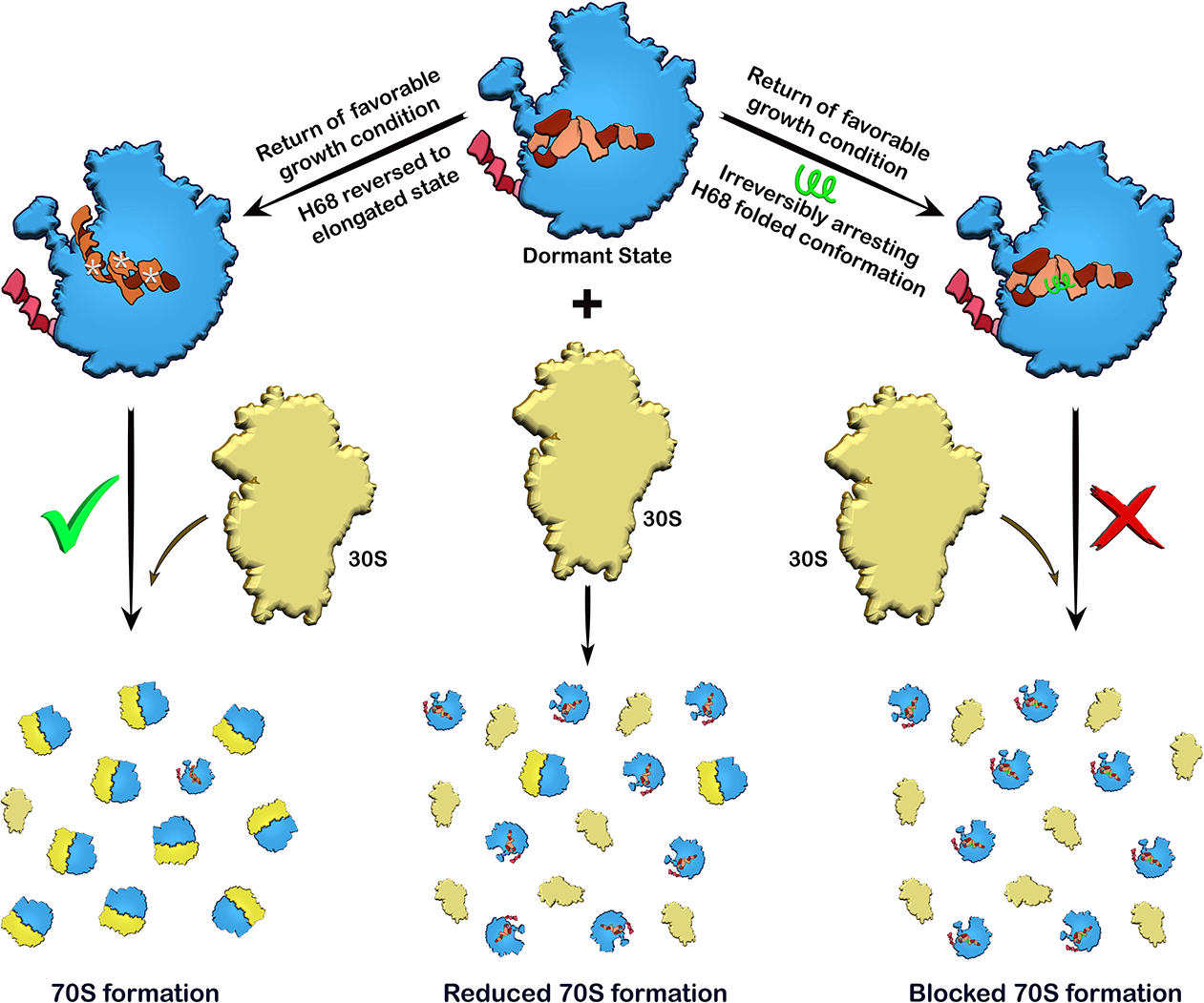
Schematic representation of implications of the conformational changes. The 50S subunit predominately acquires a folded conformation of H68 in stationary phase (mimicking dormant state) of mycobacterium. The dynamic change of H68 to folded conformation creates a pool of inactive 50S subunits in which formation of intersubunit bridges between 50S and 30S subunit is obstructed and thereby 70S formation is hindered leading to down regulation of protein synthesis (middle panel). Return to favorable growth conditions results in reversing back of H68 to elongated conformation in the 50S subunits, which makes H68, H69 and H71 accessible for intersubunit bridge formation with the 30S subunits to form active 70S ribosomes (left panel). If H68 in dormant state can be irreversibly locked in the folded conformation it would arrest protein synthesis even when bacterial growth conditions become favorable due to the scarcity of active 70S ribosome (right panel) resulting in cell death.

Alternative mycobacterial ribosomes, having distinct translational features, contribute to environmental adaptation (Chen et al., 2020). It has been shown that significant changes in the composition of ribosomal proteins occur as paralog counterpart of some ribosomal proteins get incorporated in the ribosome when cells move from exponential to stationary growth phase and from low to high Zn^2+^ medium for growth (Dow and Prisic, 2018; Lilleorg et al., 2019). Moreover, posttranscriptional modifications in H92 are well known (Hansen et al., 2002) and it has been reported that lack of U2552 (in Msm U2776) of A-loop (H92 of 23S rRNA) methylation affects the stability of H68-H71 and rRNA modification has been predicted to facilitate the conformational rearrangement of rRNA (Wang et al., 2020). Interestingly, interaction of H71 with the A-loop is rearranged along with the conformational switch in H68. Further, methylation of C2144 of H69 is also reported in a recent structural study of mycobacterial large ribosomal subunit showing that reported methylation causes flipping out of nucleotide base which is required for binding of anti-mycobacterial antibiotic capreomycin (Laughlin et al., 2022). We speculate that either a growth phase-specific rRNA modification or alteration of ribosomal protein composition might play a key role in stabilizing the altered conformation of the domain IV helices during stationary phase. It could also be possible that a yet-unknown, non-canonical factor stabilizes the dynamics of the H68 folded conformation during full dormancy.

*Mtb* in dormant state is less susceptible to antibiotics and becomes intrinsically resistant to many antibiotics including two important TB-drugs, rifampicin and isoniazide, restricting the number of available therapeutic drug molecules (Gygli et al., 2017; Luthra et al., 2018; Wayne and Sramek, 1994; Zhang et al., 2022). Our study not only provides fundamental insight into one of the mechanisms of biological adaptation of mycobacteria in translational regulation, but it also identifies a novel target for developing antimycobacterial agents. As observed in our study, H68 prefers to adopt a folded conformation in the stationary state (closely mimicking dormancy). Irreversibly arresting H68 in the folded conformation during dormant state would lead to translation shutdown in mycobacteria due to unavailability of active 70S ribosome for protein synthesis, even when the growth condition becomes favorable which ultimately results in cell death (Fig. 7).

## Acknowledgements

This work was primarily supported by SERB, DST (India) sponsored projects (SPF/2021/000141 and CRG/2019/001788), CSIR Niche Creating Project (NCP) MLP-139 and CSIR-Indian Institute of Chemical Biology, Kolkata, India. We greatly acknowledge the National Electron Cryo-Microscopy facility at the Bangalore Life Sciences Cluster (DBT/PR12422/MED /31/287/2014), NCBS, Bangalore, the Supercomputing Facility provided by CSIR Fourth Paradigm Institute (http://www.cmmacs.ernet.in), SERB, DBT (grant no. SB/SO/BB-0025/2014 and BT/PR15017/BRB/10/1445), and the Central Instrumentation Facility (CIF) of CSIR-Indian Institute of Chemical biology. We sincerely thank Dr. Vinothkumar Kutti Ragunath for helping in grid preparation and high-resolution single particle data collection. We highly appreciate Dr. Ilic Zoran’s (Wadsworth Center, NY, USA) help to refine the presentation of our manuscript. PB acknowledges CSIR, India, for research fellowship.

## Author contributions

JS conceived the project. JS, PB designed the experiments. PB carried out all the experiments including cryo-EM 3D image processing and prepared the illustrations. JS and PB analyzed the data and wrote the paper.

## Accession numbers

Cryo-EM density maps have been deposited in the Electron Microscopy Data Bank under the accession numbers EMD-33096 and EMD-33599 for *Msm* S-50S and *Msm* L1-50S, respectively. *Msm* L3-50S has been submitted as associated map to *Msm* L1-50S. The atomic models of the *Msm* S-50S and *Msm* L1-50S have been deposited in the Protein Data Bank under the accession number 7XAM and 7Y41, respectively.

## Supplementary Information

**Figure S1:**
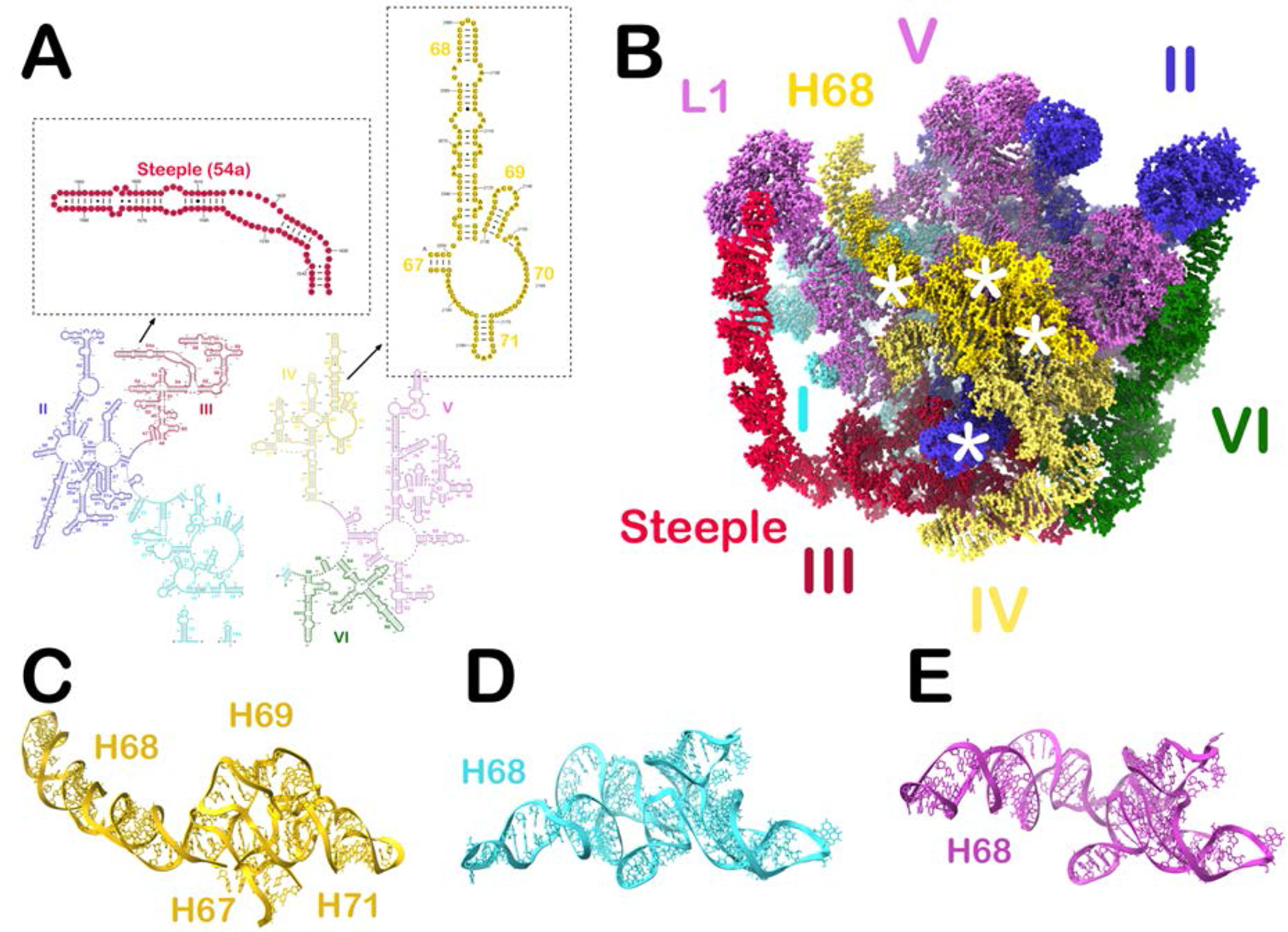
The secondary and tertiary structures of Mycobacterial 23S rRNA. (A) Secondary structures of 5’ (left), 3’ (right) ends of 23S rRNA, H54a (Steeple) and H67-H71 (domain IV) are shown in insets. (B) Colour-coded 3D folding of the domains of Mycobacterium 23S rRNA. The 50S subunit counterparts of the inter-subunit bridge forming regions are marked with white asterisks. (C) View of the H67-H71 as observed in published structures of 70S ribosome and 50S subunit of mycobacteria. RNAComposer web server predicted rotated conformation for *Msm* H68 and *Mtb* H68 (D & E), respectively.

**Figure S2:**
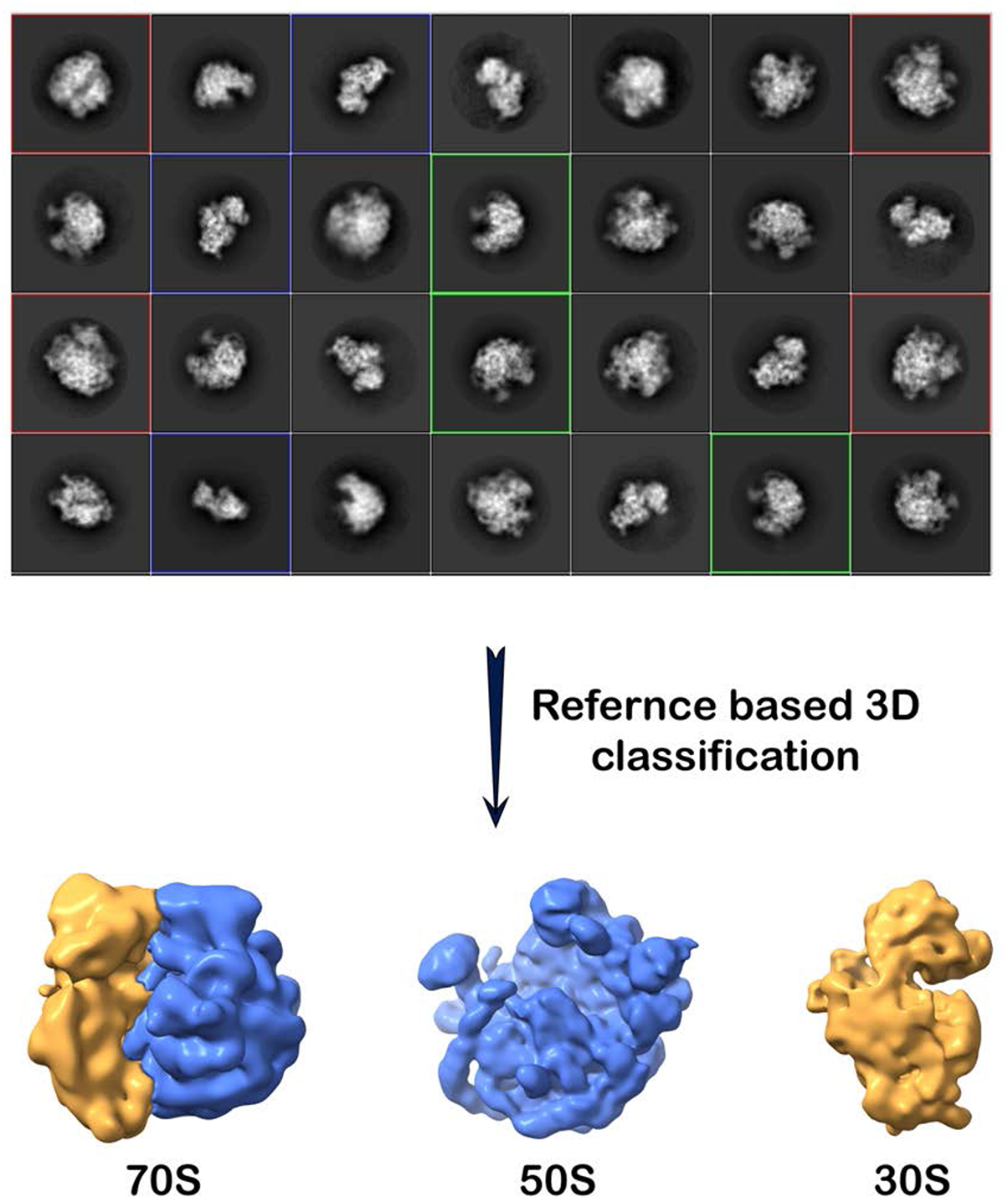
The workflow for computationally processing structural datasets in Relion. Representative views of the 2D classes of a dataset are shown (upper panel). The presence of 70S ribosome (red box), 50S (green box) and 30S subunits (blue box) is clearly visible (multiple rounds of 2D classification were done). Selected good 2D class averages were subjected to reference-based 3D classification (multiple rounds). 3D classification of the cryo-EM data processing resulted in 70S ribosome, 50S and 30S subunits density maps (lower panel). The same approach was taken to process both the log- and stationary phase datasets. Multiple rounds of 2D and 3D classifications were performed to extract the 50S subunits of log- and stationary phase ribosome datasets.

**Figure S3:**
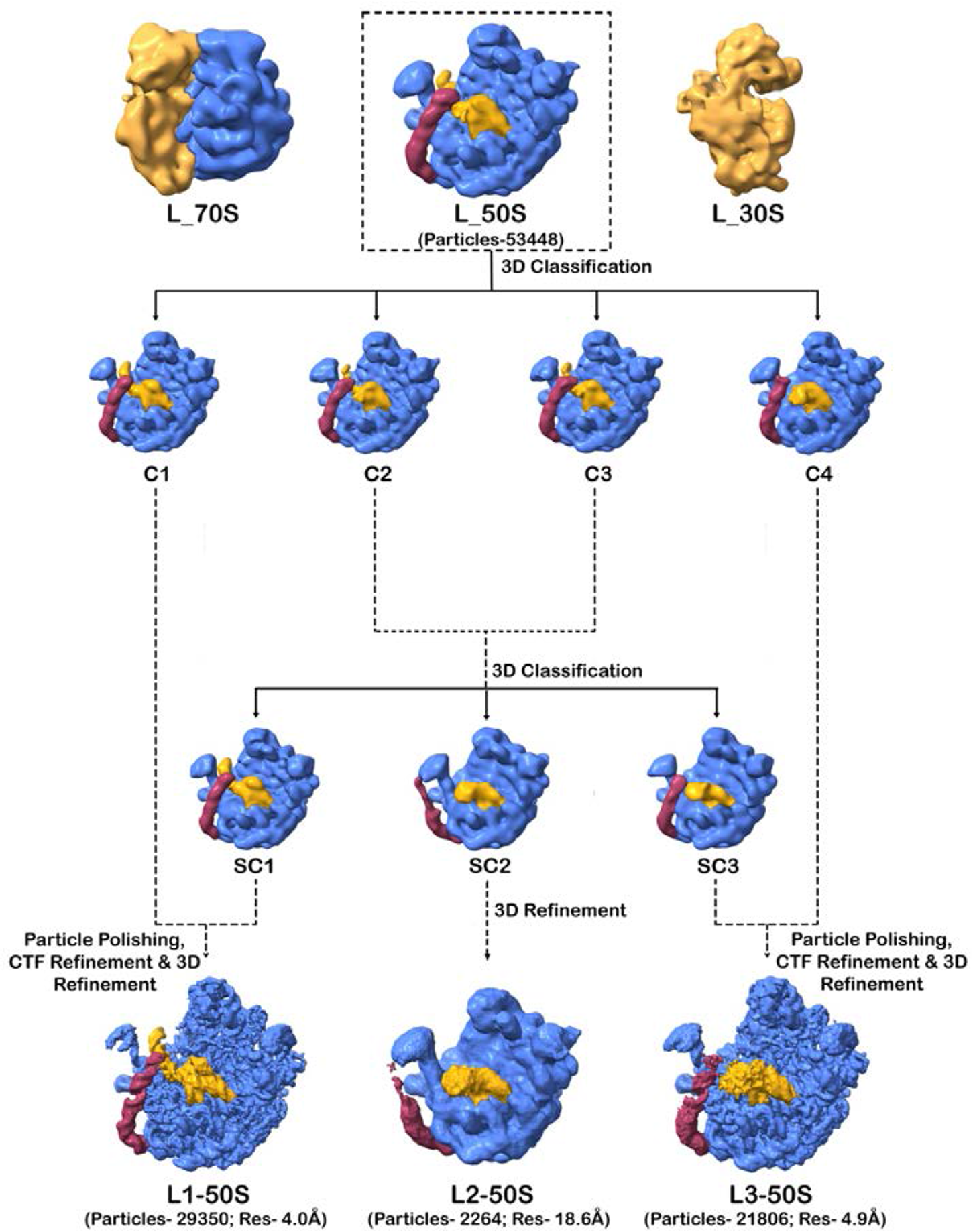
Schematic representation of 3D classification of the *Msm* 50S subunit from a log phase ribosome dataset. 3D classification of a dataset of *M. smegmatis* log phase ribosome resulted in generation of classes of 70S, 50S and 30S subunits. A class of 50S (L_50S) consisting of 53,448 particles was extracted and given for another round of reference-based 3D classification to yield 4 classes of 50S (Class 1 (C1) with unrotated H68, class 2 & 3 (C2 & C3) with mixed conformation of H68 and class 4 (C4) with rotated H68). Class 2 and 3 were merged and subjected to multiple rounds of reference-based 3D classification to get the homogenous classes. After multiple rounds of 3D classification, 3 different types of sub-classes were generated (Sub-class 1 (SC1) with unrotated H68, sub-class 2 (SC2; 2264 particles) with rotated H68 and steeple out and sub-class 3 (SC3) with rotated H68). SC1 and C1 were merged to get a total of 29,350 particles of unrotated H68, whereas SC3 and C4 were merged to get a total of 21,806 particles of rotated H68. SC2 was refined to get map L2-50S. Each particle sets of unrotated/elongated H68 and rotated H68 were subjected to particle polishing, CTF refinement and 3D refinement to generate high resolution map referred as L1-50S and L3-50S, respectively.

**Figure S4:**
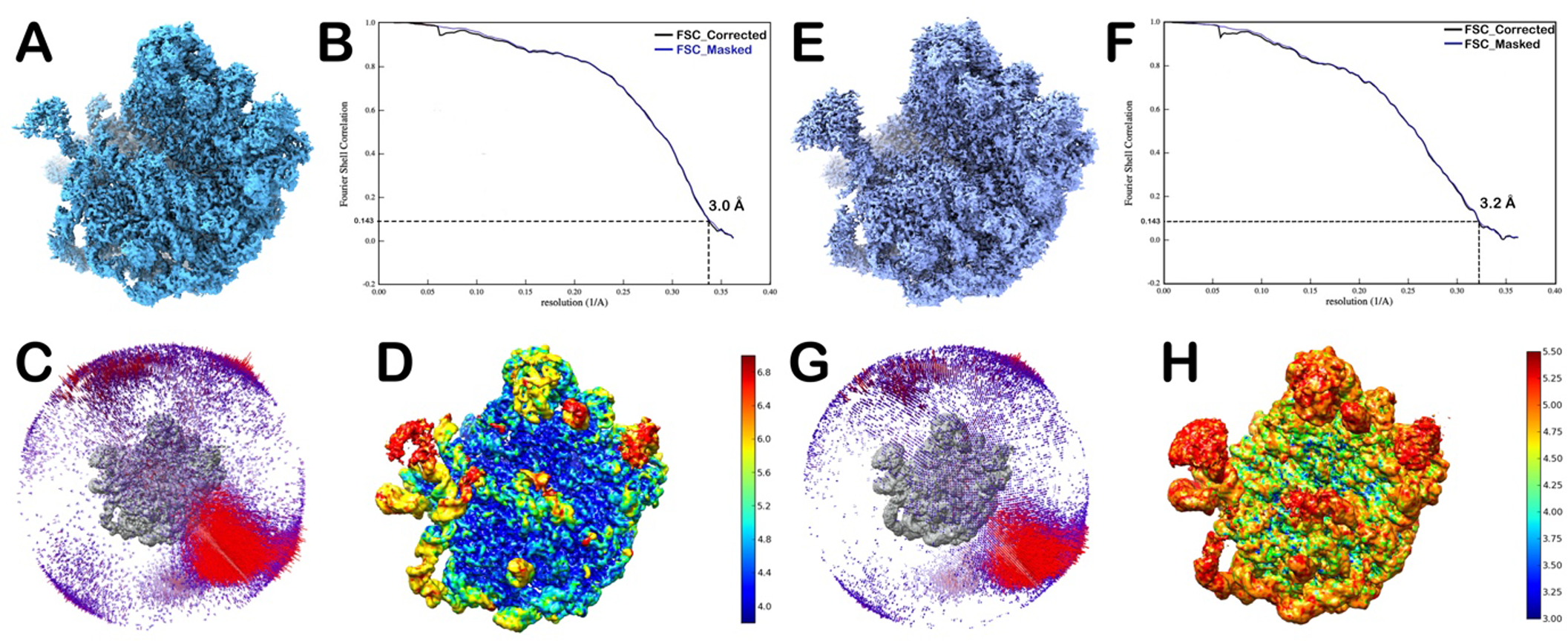
Resolution of the single particle cryo-EM reconstructions of L1-50S and L3-50S. (A & E) B-factor sharpened maps of L1-50S and L3-50. (B & F) The Fourier shell correlation (FSC) plots of L1-50S and L3-50S. Resolution values are given according to the “gold standard” (FSC = 0.143) criterion. (C & G): Angular distributions of particles of L1-50S and L3-50S datasets. (D & H): Local resolution plots of L1-50S and L3-50S generated using ResMap.

**Figure S5:**
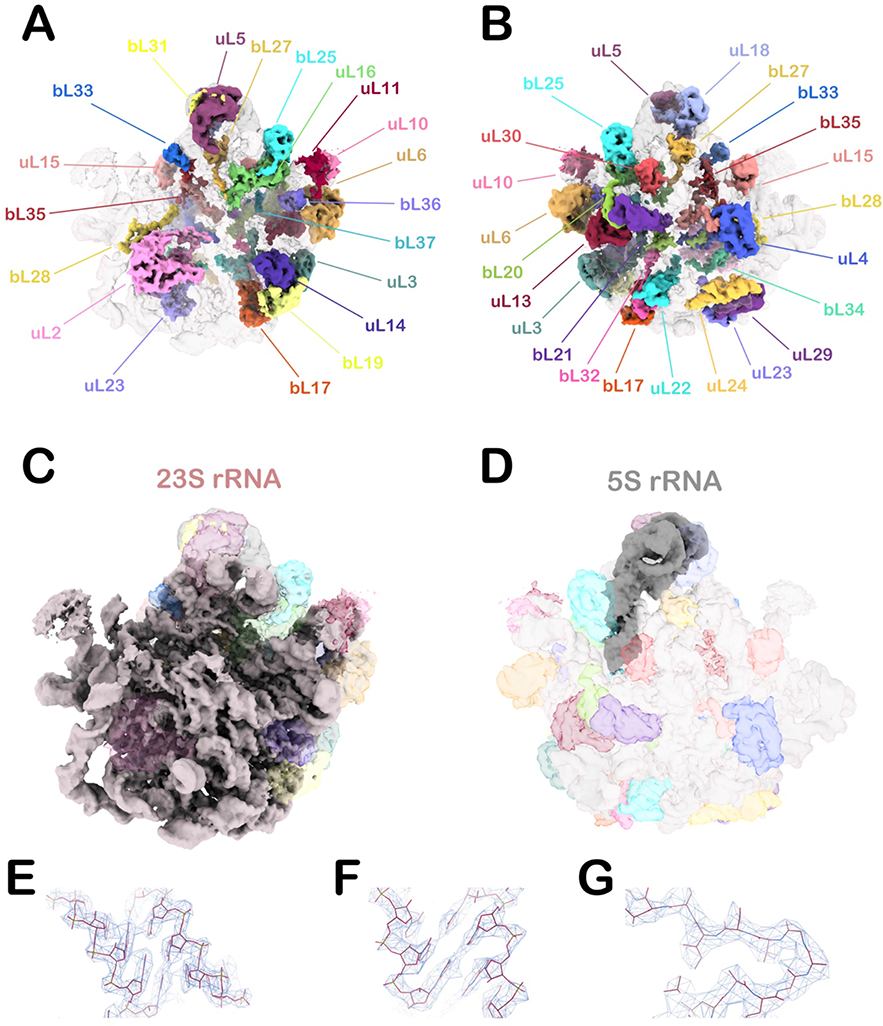
Segmentation of L1-50S cryo-EM map and fitted atomic model. (A & B) Front and back view of segmented ribosomal proteins from L1-50S. Full densities corresponding to all the ribosomal proteins (except uL9) are present (colour-coded). (C) Intersubunit view of segmented density of 23S rRNA, and (D) segmented density of 5S rRNA from back view of 50S subunit are provided. Atomic model fitting of rRNA (E-F) and ribosomal protein (G) in representative segments of cryo-EM density map of L1-50S are shown.

**Figure S6:**
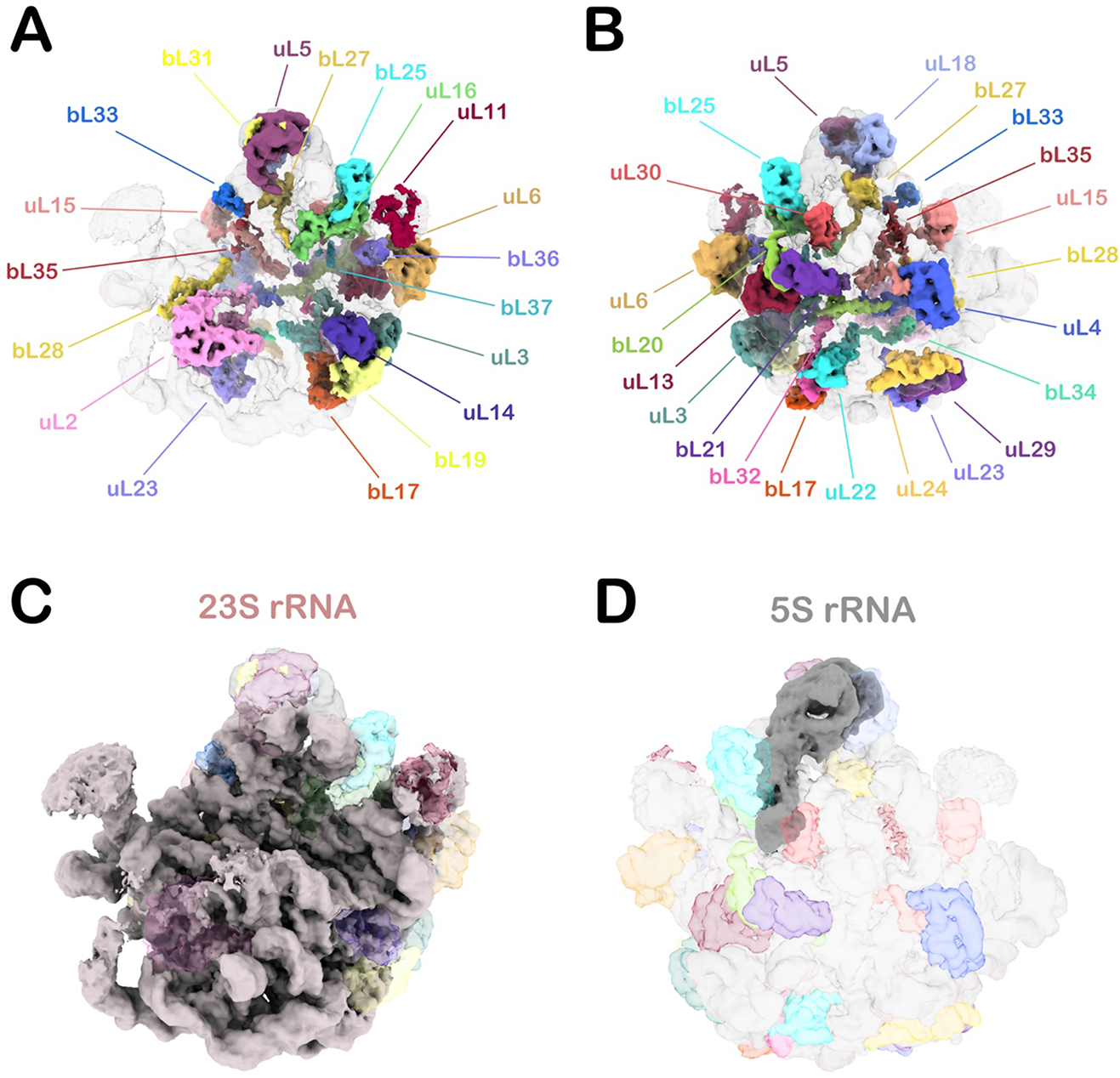
Segmentation of L3-50S cryo-EM map. (A & B) Front and back view of segmented ribosomal proteins from L3-50S. Full densities corresponding to all the ribosomal proteins (except uL9 and uL10) are present (colour-coded). (C) Intersubunit view of segmented density of 23S rRNA, and (D) segmented density of 5S rRNA from back view of 50S subunit are given.

**Figure S7:**
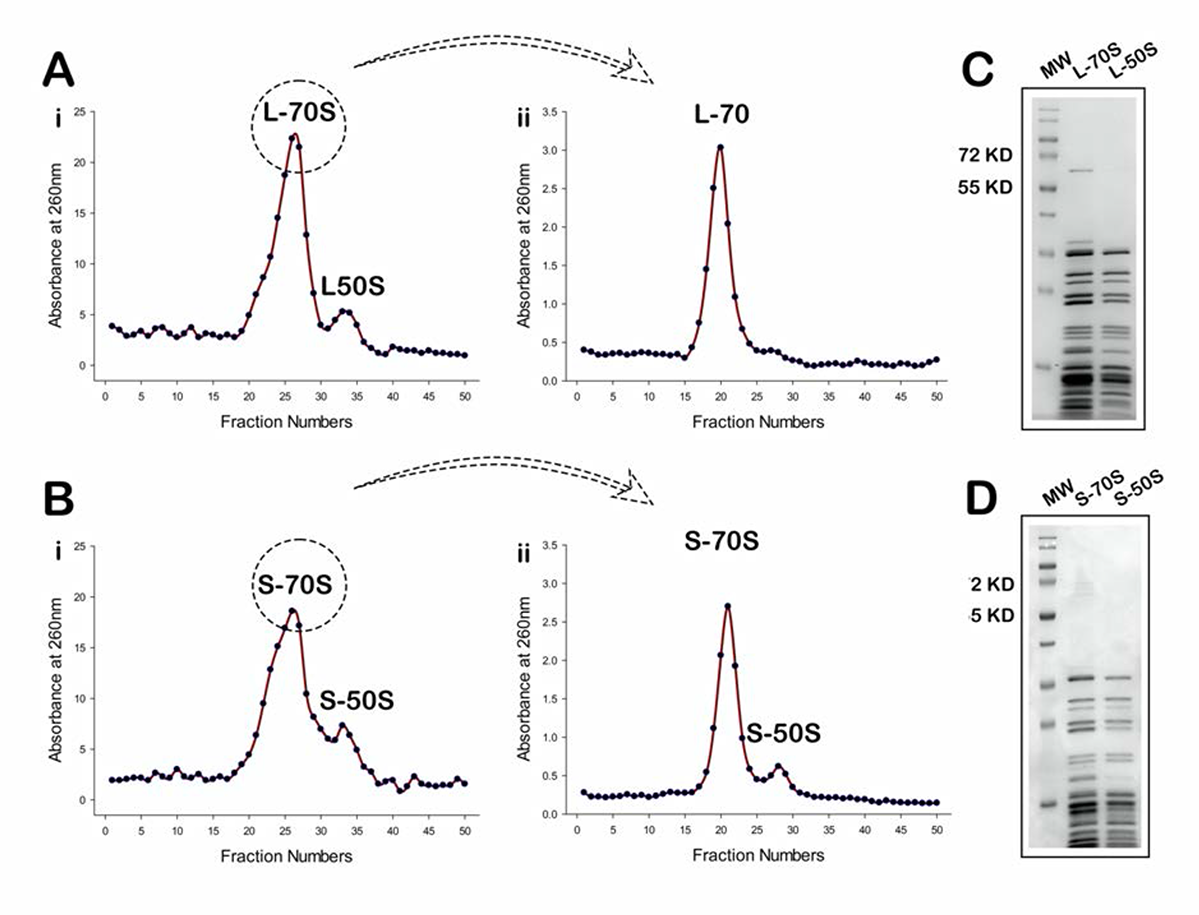
Analysis of ribosome profile of *M. smegmatis* log and stationary phase ribosomes. (A & B) Sucrose density gradient (SDG) ribosome profile of purified 70S ribosome from log phase (L-70S) (A: left panel) and stationary phase (S-70S) (B: left panel). Stored 70S (L-70S & S-70S) were reloaded onto sucrose density gradient. SDG ribosome profile shows L-70S retained 70S ribosome pool (A: right panel), whereas, one additional peak of S-50S is seen along with S-70S (B: right panel) suggesting self-dissociation of S70S creates an inactive pool of free S-50S ribosomal subunits of stationary phase causing equilibrium shift, which further prevents re-association with 30S ribosomal subunit to form active ribosome. (C & D) SDS PAGE profiles show the signature bands of ribosomal proteins of purified 70S ribosome and 50S subunit from log and stationary phases, respectively. Purification of ribosome was done multiple times. One of the representative profile is shown over here.

**Figure S8:**
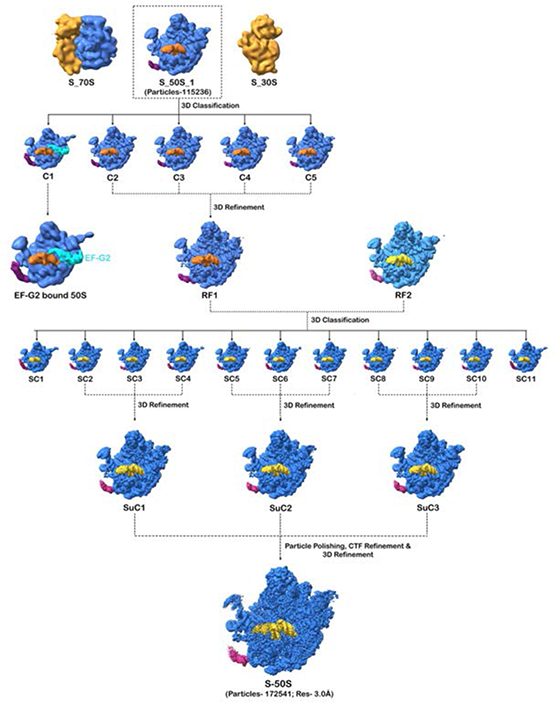
Schematic representation of 3D classification of the *Msm* 50S subunit from stationary phase ribosome dataset. 3D classification of a dataset of *M. smegmatis* stationary phase ribosome resulted in generation of classes of 70S, 50S and 30S subunits. The 50S subunit class of dataset one (S_50S_1) consisted of 115236 particles, subjected to a series of 2D averaging and 3D classification. Finally, five classes of 50S were generated, out of which one class was EF-G bound (C1) having ∼ 5,700 particles, whereas all other classes showed the same 3D conformation with rotated H68 and steeple out (C2-C5; total consisting of 1,04,076 particles). The four classes showing the same conformation were combined and subjected to 3D refinement to generate map RF1. 50S refined map (RF2) from another *Msm* S-ribosome-EF-G2 complex dataset having the same conformation were combined with 50S refined map (RF1) of *Msm* S-ribosome-EF-G2-GDPNP complex dataset. Combined 50S particles were subjected to a series of 2D classification to remove some bad particles. Finally, good 2D class averages were subjected to reference-based 3D classification, resulting in formation of 11 major 3D Classes (SC1-SC11), out of which 9 classes were found to be identical (rotated H68 and steeple out), and further merged as set of 3-3 classes to generate 3 Super classes (SuC1-SuC3). All three super classes were combined to generate a final refined map with 1,72,541 particles. 3D refined map was subjected to particle polishing, CTF refinement and 3D refinement to generate high resolution map named as S-50S.

**Figure S9:**
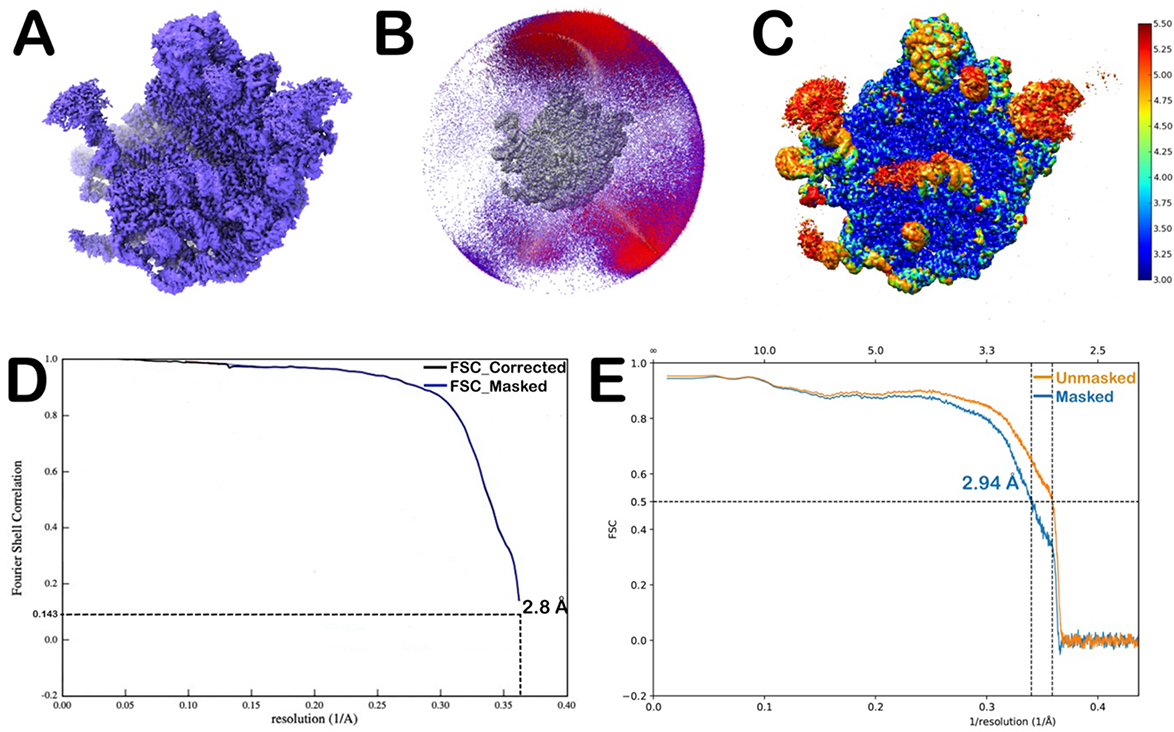
Resolution of the single particle cryo-EM reconstructions and model of S-50S. (A) B-factor sharpened maps of S-50S. (B) Angular distribution of particles of S-50S. (C) Local resolution plot of S-50S was generated using ResMap (13). (D & E) The Fourier shell correlation (FSC) plots of the map (according to the “gold standard” FSC=0.143 criterion) and model of S-50S.

**Figure S10:**
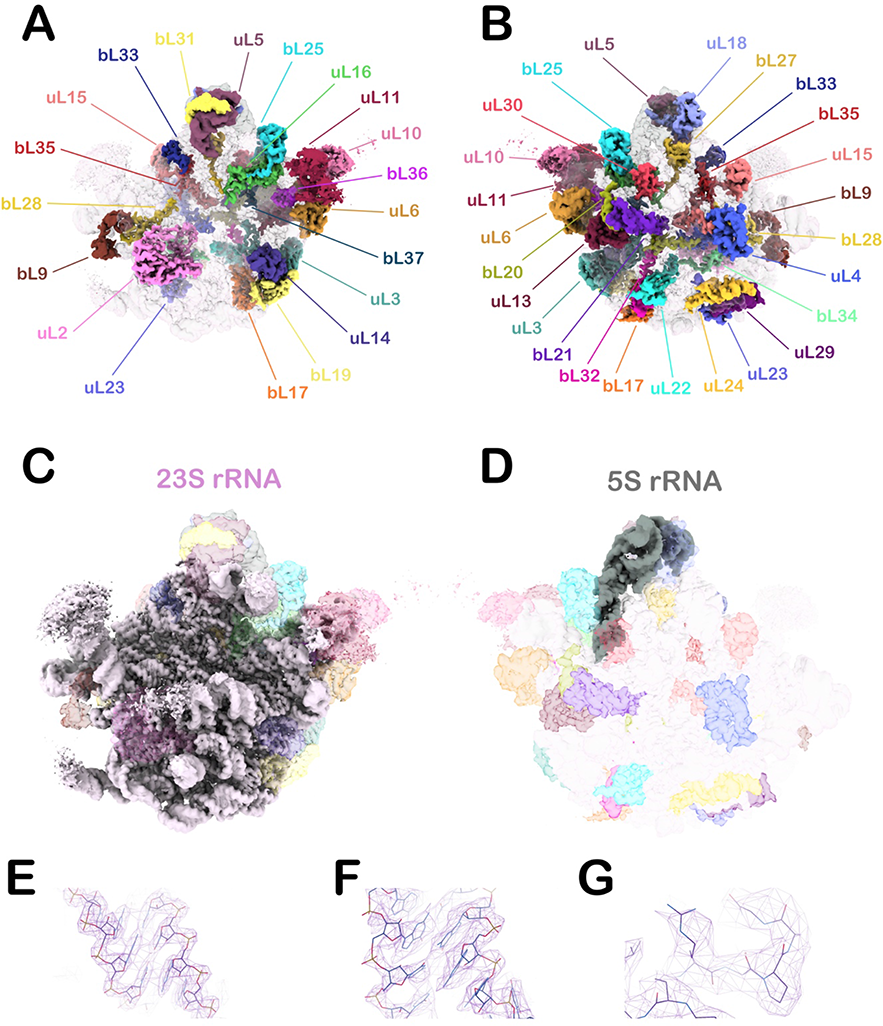
Segmentation of S-50S cryo-EM map and fitted atomic model. (A & B) Front and back view of segmented ribosomal proteins from S-50S. Full densities corresponding to all the ribosomal proteins (colour-coded) are clearly visible. (C) Intersubunit view of segmented density of 23S rRNA. (D) Segmented density of 5S rRNA from back view of 50S subunit. Atomic model fitting of rRNA (E-F) and ribosomal protein (G) in representative segments of cryo-EM density map of S-50S.

**Figure S11:**
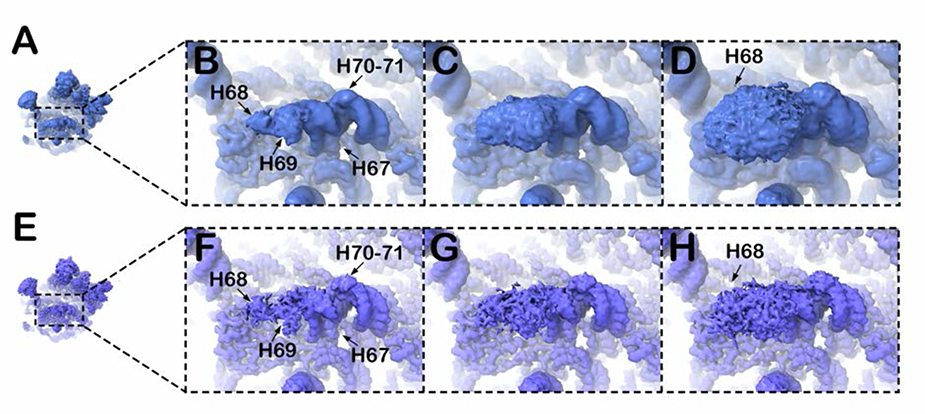
Representation of H67-H71 of S-50S in refined and sharpened maps at different contour levels. (A & E): Thumbnails of cryo-EM map of S-50S subunit following refinement (3.1Å) and following b-factor sharpening (2.8Å), respectively. (B-D & F-H) Close-up views of H67-71 region of domain IV from high to low threshold in the refined map (B-D) and in sharpened map (F-H). The density corresponding to H70-71 is seen at high threshold in both the maps, whereas, the fused density attributable to the folded H68-H69 is weak. The mushroom head-like density is visualized in the refined map at lower threshold and allowed us to build the backbone trace model of this region. This density, however, appears fragmented in the sharpened map suggesting conformational variability of this region.

**Figure S12:**
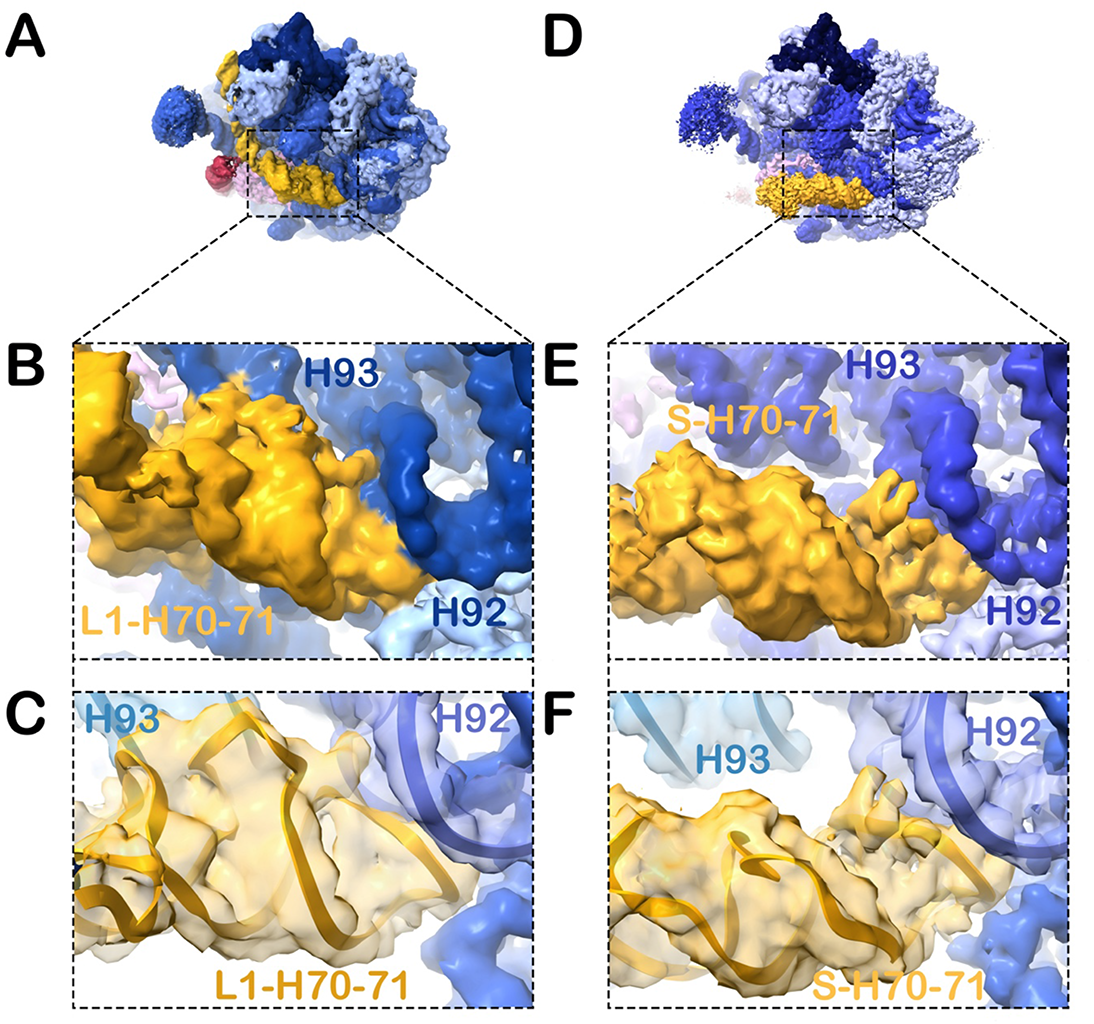
Comparison of H70-H71 conformational changes in L1-50S and S-50S. (A & D): Top views of the cryo-EM density maps of L1-50S and S-50S. (B-C): Molecular arrangement of partly unstructured H70-71 in L1-50S shows its interactions with nearby H92 and H93 of domain V as visualized in density map and molecular model. (E-F): Structural rearrangement of H7-71 is seen in density map and model of S-50S due to the breakage of molecular interactions with H93 which results in its forward movement towards inter-subunit space.

**Figure S13:**
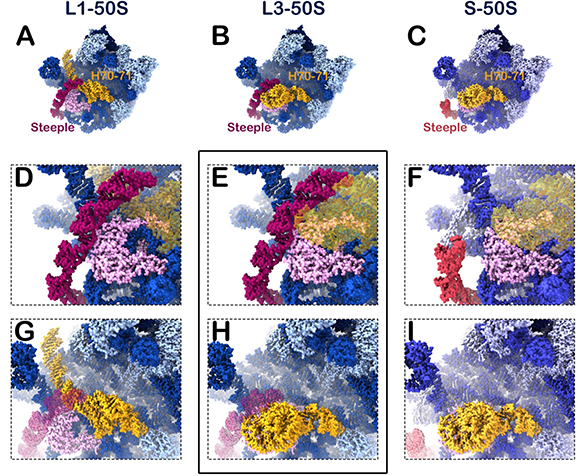
Comparative analysis of Steeple (H54a) and H67-H71 in L1-50S, L3-50S and S-50S. (A-C) Intersubunit views of the molecular models of L1-50S, L3-50S and S-50S. (D-F) Steeple (H54a) stays in tilted position in L1-50S and L3-50S, whereas, it moves outward in S-50S. (G-I) Elongated structure of H68 of domain IV is seen in L1-50S, while it attains folded conformation in L3-50S and S-50S. It appears from the comparative analysis of the positions of Steeple and H67-H71 that L3-50S represents an intermediate conformation between L1-50S and S-50S. H54a seems to be destabilized when its interaction with H68 in elongated conformation gets disrupted.

## Movie legends

**Movie S1: Drastic conformational switching of H68.** The elongated conformation of H68 in L1-50S transforms to a folded conformation in S-50S.

**Movie S2: Dynamic behaviour of H67.** H67 slightly tilts towards intersubunit space (∼5-6 Å) in S-50S from its position in L1-50S.

**Movie S3: Structural rearrangement of H70-H71.** Significant intra-helical conformational rearrangement is observed in H70-H71 upon transition from L1-50S state to S-50S state.

**Movie S4: Conformational transition of H69.** H69 flips laterally ∼90° in S-50S as compared to its position in L1-50S.

**Movie S5: Anti-correlated motions of H68 and Steeple (H54a)**. Comparison of L1-50S and S-50S structures manifests that motions of H68 and Steeple (H54a) of the *Msm* 23S rRNA are in opposite directions.

**Note:** All movies are prepared using ChimeraX and online GIF maker.

